# An EMG-assisted Muscle-Force Driven Finite Element Analysis Pipeline to Investigate Joint- and Tissue-Level Mechanical Responses in Functional Activities: Towards a Rapid Assessment Toolbox

**DOI:** 10.1101/2021.03.27.436509

**Authors:** A. Esrafilian, L. Stenroth, M. E. Mononen, P. Vartiainen, P. Tanska, P. A. Karjalainen, J. S. Suomalainen, J. Arokoski, D. G. Lloyd, R. K. Korhonen

## Abstract

Joint tissue mechanics (e.g., stress and strain) are believed to have a major involvement in the onset and progression of musculoskeletal disorders, e.g., knee osteoarthritis (KOA). Accordingly, considerable efforts have been made to develop musculoskeletal finite element (MS-FE) models to estimate highly-detailed tissue mechanics that predict cartilage degeneration. However, creating such models is time-consuming and requires advanced expertise. This limits these complex, yet promising MS-FE models to research applications with few participants and making the models impractical for clinical assessments. Also, these previously developed MS-FE models are not assessed for any activities other than the gait. This study introduces and validates a semi-automated rapid state-of-the-art MS-FE modeling and simulation toolbox incorporating an electromyography (EMG) assisted MS model and a muscle-force driven FE model of the knee with fibril-reinforced poro(visco)elastic cartilages and menisci. To showcase the usability of the pipeline, we estimated joint- and tissue-level knee mechanics in 15 KOA individuals performing different daily activities. The pipeline was validated by comparing the estimated muscle activations and joint mechanics to existing experimental data. Also, to examine the importance of EMG-assisted MS analyses, results were compared against outputs from the same FE models but driven by static-optimization-based MS models. The EMG-assisted MS-FE pipeline bore a closer resemblance to experiments, compared to the static-optimization-based MS-FE pipeline. More importantly, the developed pipeline showed great potentials as a rapid MS-FE analysis toolbox to investigate multiscale knee mechanics during different activities of individuals with KOA.

## 1. Introduction

Knee injuries, after back disorders, are the second most frequent musculoskeletal (MS) disorder [1]. There is compelling evidence that altered knee joint motion and loading, and the subsequent mechanical responses (i.e., stress and strain) within the knee load-bearing tissues, are key factors in the onset and progression of knee osteoarthritis (KOA) [2–9]. Hence, a thorough knowledge of the tissue mechanical responses to knee joint loading is essential to assess KOA and possibly restore knee functionality. In this regard, knee joint contact forces (JCF), contact area, and contact pressure have been experimentally measured in several activities [4, 10–16]. These experiments have revealed fundamental information on the knee joint mechanics, but they are either limited to specific subjects (e.g., those with instrumented implants) or require highly invasive procedures. More importantly, experimental approaches cannot measure crucial parameters governing tissue adaptation and degeneration, such as stress, strain, or fluid flow of the tissue.

Alternatively, multiscale MS and finite element (FE) models have become a tool of choice to investigate detailed joint loading and tissue-level mechanical responses [6, 17–23]. However, none of the developed multiscale MS-FE models have been used for analyzing joint mechanics in functional activities other than gait. Furthermore, to the best of our knowledge, there are no studies incorporating subject-specific muscle recruitment (activation) strategies in the estimation of tissue-level mechanical responses, although muscle recruitment has a significant effect on the joint loading, especially in the presence of MS disorders [24–28]. Moreover, those studies reporting detailed joint mechanical responses (either joint-level or tissue-level) have investigated healthy subjects [17,28–30], utilized simplified joint models in terms of limited degrees of freedom (DoFs) [30–32], excluded subject-specific joint geometries [28, 30–32], omitted crucial joint tissues (e.g., menisci) [28, 30, 31], and/or utilized simple soft tissue material models [28]. Patient-specific joint geometries [33], inclusion of menisci [34–36] and a multi DoFs joint model [20], and the use of an appropriate soft-tissue material model can substantially alter the estimated tissue mechanics [37–41].

A fibril-reinforced composite material model is essential to be able to simultaneously estimate mechanical responses of both the fibrillar (collagen) and nonfibrillar (proteoglycans) matrices (e.g., in cartilage and meniscus) [37,42,43]. Moreover, poroviscoelasticity is needed to replicate fluid-flow-dependent and -independent mechanisms of biphasic tissues [44], with within-tissue fluid pressurization carrying up to 90% of a dynamic load [40, 45]. These characteristics of the knee soft tissue emphasize the use of a fibril-reinforced poroviscoelastic (FRPVE) material model, which can potentially provide the FE analysis with a more detailed estimation of tissue-level mechanical responses, especially if adaptation and degradation of cartilage and its fibrillar and nonfibrillar matrices are of interest [6, 43, 46, 47].

Summarizing, FE models utilizing subject-specific joint geometries and complex FRPVE material models have shown great potential for estimating highly-detailed tissue mechanics and predicting cartilage damage and degeneration [6, 39, 43, 46–48]. However, creating the above-mentioned FRPVE FE models is a cumbersome manual task requiring several weeks of high-level expertise [48]. This process entails image segmentation, meshing, material model incorporation, estimation and application of loading and boundary conditions, and getting a converged solution. This lengthy procedure limits these complex, yet promising, FE models to research purposes with only a few participants, and therefore their application is impractical and infeasible for large cohorts or clinical assessments. Some efforts have been made to be able to rapidly create FRPVE FE models of the knee, but they are strictly limited to the tibiofemoral joint during walking without subject-specific loading and boundary conditions [49].

Therefore, here we develop a rapid state-of-the-art MS-FE modeling and simulation pipeline potentially feasible for research and clinical applications to investigate joint- and tissue-level knee mechanics in different functional activities. To this end, we adapted an atlas-based FE modeling toolbox [49] along with an EMG-assisted muscle-force driven FRPVE FE analysis workflow [17]. To showcase the usability of the pipeline, we estimated joint- and tissue-level knee mechanics in a sample of individuals with KOA while performing different daily activities. The pipeline was validated by comparing the estimated muscle activations, JCFs, and tissue mechanical responses to existing experimental data. To explore the influence of EMG-assisted MS analyses, different joint-level and tissue-level results estimated by the developed workflow were compared with the results estimated by a similar workflow but with the FE models driven by static-optimization (SO) based MS models.

## 2. Method

### 2.1. Data collection and pre-processing

Fifteen subjects (6 males and 9 females, 62.4±7.8 years old, and with body mass index 29.3±6.8) meeting the study admission criteria participated in this study (workflow in Figs. 1 and 2). Subjects’ characteristics are provided in the supplementary material, Table S1. The inclusion criterion was previously diagnosed KOA according to the KOA clinical definition (i.e., the existence of both pain and an evident radiographic joint tissue deterioration [50]) in either of the medial or lateral femur, tibia, or patella. The exclusion criteria were the existence of any record of lower limb surgeries or diagnosed disorders such as ligament or tendon rupture or presence of pain in any body parts except for the knee. Analyses were undertaken on the most affected leg of each subject (in terms of OA severity), comprising of a total of 15 knees, one knee from each subject. All the procedures were approved by the Human Research Ethics committee of the Northern Savo Hospital District (permission number 750/2018), and written informed consent was obtained from each subject.

**Fig. 1:**
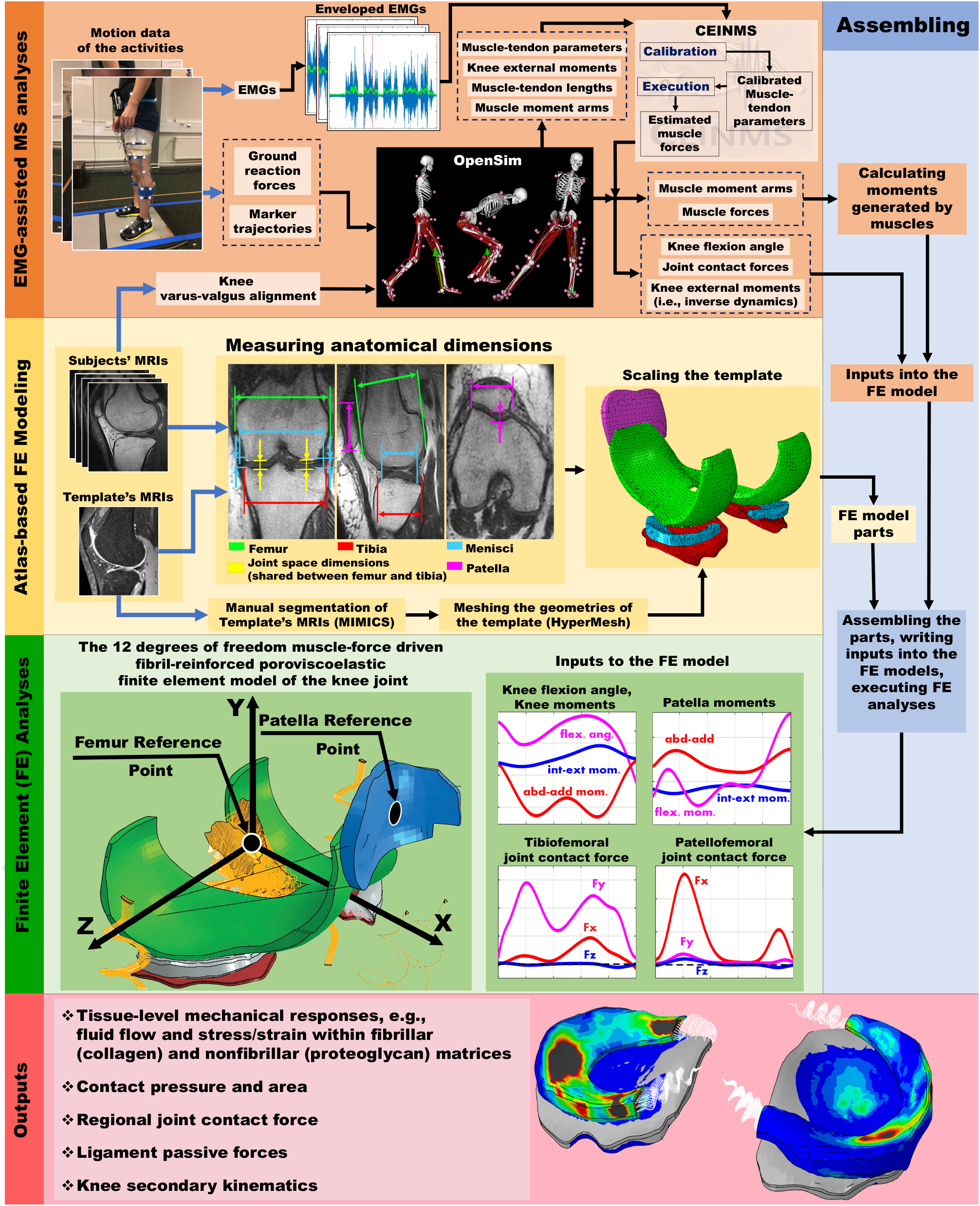
The illustration of the developed MS-FE analysis pipeline. Inputs to the pipeline are shown with blue arrows.

**Fig. 2:**
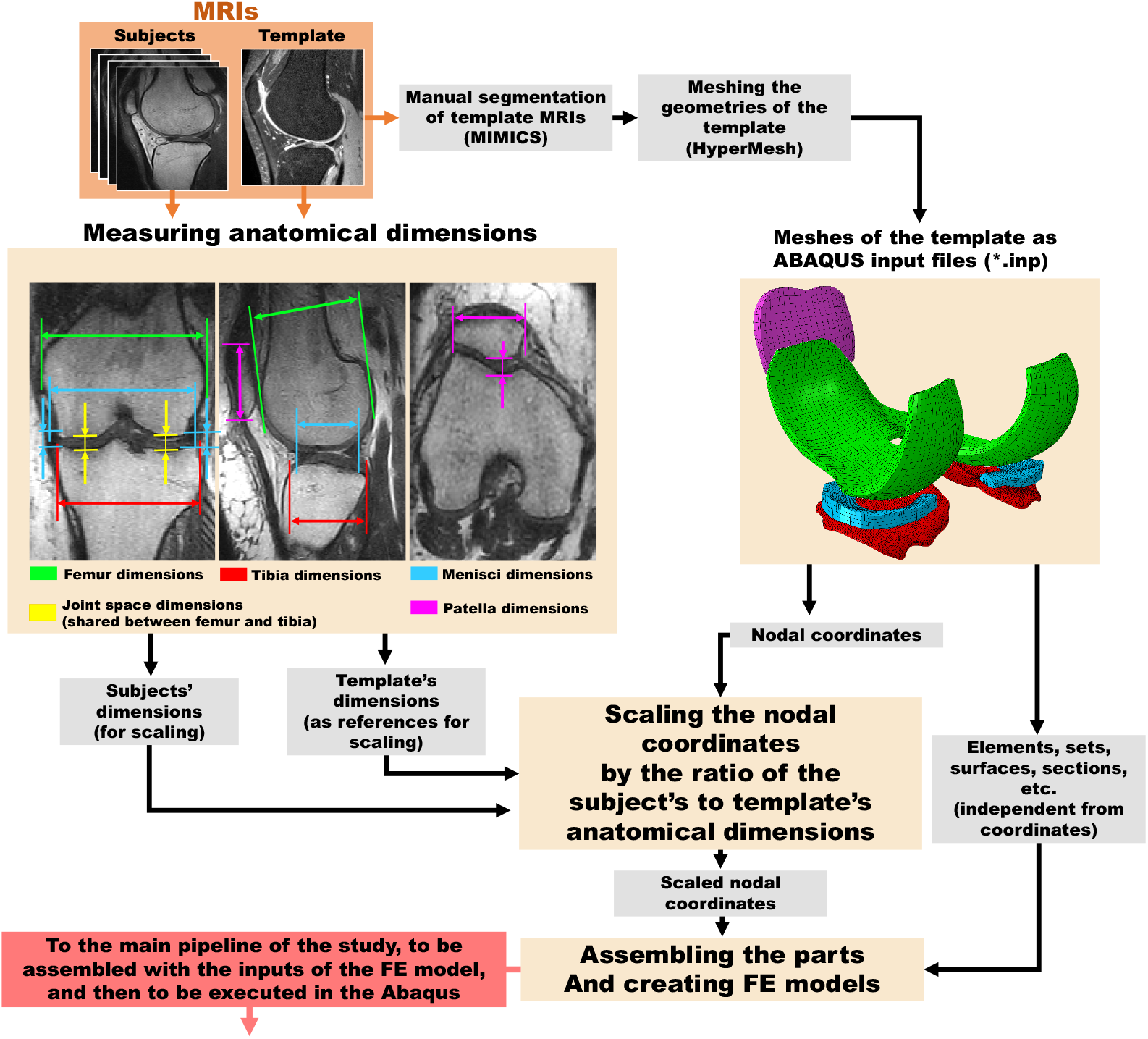
The workflow of the atlas-based FE modeling toolbox. Orange arrows show inputs to the workflow, and the light-red box shows the output of the atlas-based FE modeling toolbox to the main workflow of the study.

We analyzed seven different daily activities. These consisted of: (1) chair stand-to-sit, (2) chair sit-to-stand, (3) walking at a naturally selected speed (1.34 ± 0.14 m/s), (4) walking at a standardized speed (1.20 ± 0.05 m/s), (5) picking up a pen from the ground, (6) stair ascent, and (7) stair descent. The motion data collection (Fig. 1) consisted of synchronous measurement of 3D marker trajectories (100 Hz, Vicon, UK), ground reaction forces (two force plates, 1000 Hz, OR6-7MA, AMTI, USA), and EMGs (1000 Hz, ME6000, Bittium Biosignals Ltd, Finland). EMGs from 8 muscles of the test leg, comprising the vastus medialis and lateralis, rectus femoris, medial and lateral gastrocnemius, biceps femoris, semitendinosus and gluteus medius, were recorded according to the SENIAM guidelines [51]. Additionally, magnetic resonance images (MRIs) were taken from each subject’s test knee (Fig. 1) using a 0.18 T scanner (3D CE sequence, with 0.89 mm slice thickness and 0.625 in-plane resolution, Esaote E-Scan XQ, Esaote, Genoa, Italy). The 3D CE sequence of the utilized MRI scanner was earlier shown to be adequate to segment knee joint cartialge with a similar volume and thickness, compared to a 3T scanner (Philips, Best, Netherlands) [6].

Marker trajectories and GRFs were filtered using a fourth-order zero-lag Butterworth low-pass filter with cut-off frequencies of 6 Hz and 30 Hz, respectively. Employing MOtoNMS [52], EMG envelopes (Figs. S1 to S7) were generated from the recorded EMG signals by band-pass filtering(30-300 Hz), full-wave rectifying, low-pass filtering (6 Hz), and then normalizing to the peak similarly-processed EMG data recorded from maximum isometric voluntary exertion trials or daily activities trials the subject undertook [53]. The maximum isometric voluntary exertion trials were conducted for hip abduction, hip flexion, knee flexion/extension, and ankle plantar flexion.

### 2.2. The MS analyses pipeline

#### 2.2.1. The MS model and inputs to the MS analyses

The MS analyses consisted of the static-optimization (SO) and EMG-assisted neural solutions using an MS model with a 1 DoF knee joint optimized for modeling activities with deep knee and hip flexions [54]. The abduction/adduction and internal/external degrees of freedom were added to the tibiofemoral joint mechanism, but locked during analyses, to calculate knee abduction/adduction and internal/external moments and muscle moment arms. Similarly, flexion/extension, abduction/adduction, and internal/external DoFs were added, but also locked, to the patellofemoral joint to be able to calculate muscle moment arms around the patella. Utilizing OpenSim (v 4.1) [55], the body segments and length-dependent muscle properties of the MS model were first scaled for each subject, using the body mass and the static trial of the subject. During the scaling process, the tibiofemoral abduction-adduction DoF was opened. Then, the locations of the virtual markers around the knee were adjusted according to the MRIs, allowing the adduction angle to be adjusted subject-specifically. After scaling, the knee adduction DoF was locked. Finally, the maximum isometric muscle forces were scaled by the ratio of the subject’s mass to the mass of the un-scaled model.

Within OpenSim, the scaled models (i.e., 15 MS models in total) were used to calculate inverse kinematics (inverse kinematics toolbox), knee external moments (inverse dynamics toolbox), JCFs (for both tibiofemoral and patellofemoral joint within the joint reaction analyses toolbox), and muscle moment arms and muscle-tendon lengths (muscle analysis toolbox). The muscle moment arms were extracted for flexion/extension, abduction/adduction, and internal/external DoFs of both the tibiofemoral and patellofemoral joints. The variables were fed into the SO-based and EMG-assisted MS analyses, and then into the FE models of the study, correspondingly (Fig.1 and sections 2.2.2 and 2.3.2).

#### 2.2.2. The static-optimization and EMG-assisted MS analyses

Both the SO-based and EMG-assisted MS analyses were used to drive the FE models to investigate possible alterations in the joint-level and tissue-level mechanics from the different neural solutions. The OpenSim static optimization (SO) toolbox was used for the SO-based estimation of muscle activation patterns and forces. In this, muscle forces were estimated to track the joint moments while minimizing the sum of squared muscle activations. Muscle contraction dynamics was included, but the muscle activation dynamics was not considered within the SO analyses of OpenSim [55].

The EMG-assisted estimation of muscle activation patterns and forces was performed using the Calibrated EMG-Informed Neuromusculoskeletal Modelling Toolbox (CEINMS) [17, 22]. Inputs to CEINMS consisted of 1) muscle properties, 2) enveloped EMGs, 3) joint external moments of the leg of interest (OpenSim inverse dynamics), and 4) muscles’ moment arms and muscle-tendon lengths. Muscle properties of all the 43 muscles of the leg of interest were imported to CEINMS, including maximum isometric force, tendon slack length, optimal fiber length, and pennation angle of the muscles, which were obtained from the scaled MS models of the study, separately for each subject.

Within CEINMS, first a multi-DoFs calibration [17, 22, 56] was performed to optimize the neuromuscular parameters of all the 40 muscles of the leg of interest, separately for each subject. Five DoFs were included, consisted of the hip (3 DoFs), knee (1 DoF), and ankle (1 DoF). The neuromuscular parameters were the maximum isometric force, tendon slack-length, optimal fiber-length, EMG-to-activation recursive filter-coefficients, and nonlinear shape-factor [53, 57]. One trial of each daily task was included in the calibration. Following calibration, the hybrid mode of the CEINMS toolbox, with elastic tendons and including muscle activation and contraction dynamics, was employed to perform the EMG-assisted MS analyses (Fig. 1). In this, a simulated annealing algorithm was used to minimize the following cost function:

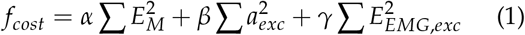

where *E_M_* is the error between joint moment estimated by the inverse dynamics and the joint moment generated by the estimated muscle forces at each DoF, *a_exc_* is the estimated excitation of each of the 40 muscles, and *E_EMG_*,*_exc_* is the error between the EMG envelopes and the corresponding estimated muscle excitations. The weight factors *α* and *β* were set to one, while the *γ* was obtained (relative to *α* and *β*) through optimization to ensure equally minimized joint moment and muscle excitation errors [58].

The muscle forces from both neural solution approaches were fed into the OpenSim’s joint reaction analysis toolbox to calculate the JCFs for all the activities. Subsequently, both SO-based and EMG-assisted estimated JCF, joint moments, muscle forces, and muscle moment arms were used to drive the FE model (Figs. 1 and section 2.3.2).

### 2.3. FE analyses pipeline

#### 2.3.1. The atlas-based FE modeling toolbox

To develop a workflow for the rapid generation and simulation of the MS-FE model, we used a novel atlas-based FE modeling approach [49] along with a muscle-force driven FE analysis workflow [17]. It was earlier shown that tissue mechanics estimated by using the one template approach, i.e., compared to choosing between several templates, could better predict KOA progression and classify subjects into correct KOA grade groups consistent with experiments [49].

Hence, we used the geometries of the FE model from our previous study [17, 23] as the template. Knee cartilages, menisci, and ligament insertion points were manually segmented (MIMICS, version 21, Materialise, Belgium) and the 3D geometries were meshed precisely in HyperMesh (version 2019, Altair, US). All the meshed geometries were then imported into the Abaqus software (version 6.20, Dassault Systèmes, US) to create a complete FE model of the template subject. All the parts were assembled, and ligament bundles and menisci horn attachments were defined according to the insertion points obtained from the template MRIs. The reference points (Fig. 1), node sets, and element sets required for applying boundaries and loads, contacts, and material models, as well as those sets for reading the results, were defined for each part. Next, material models were assigned, and contacts and couplings were defined. Femoral, tibial, and patellar cartilage were modeled utilizing the FRPVE material model [59–61, 61–66], and menisci were modeled as a fibril-reinforced poroelastic material [67]. Knee ligaments, including anterior and posterior cruciate ligaments, medial and lateral collateral ligaments, lateral and medial patellofemoral ligament, patellar tendon, and menisci horn attachments, were modeled as spring bundles [68–71].

Finally, the whole FE model of the template was exported as an Abaqus input file (.inp extension). The generated template was then scaled in a patient-specific manner according to the morphological dimensions of each subject [49] (Fig. 2). The morphometry used for scaling the template were measured from MRIs and consisted of mediolateral, anteroposterior, and transverse dimensions, or alternatively, thicknesses (to scale menisci) and joint spaces (to scale femoral, tibial, and patellar cartilage thicknesses) [49] (Fig. 2). These nodal coordinates were used to scale the template file (except for kinematic and kinetic inputs) while the rest of the Abaqus input file (i.e., element definitions, node and element sets, etc.) was identical for all the subjects (Fig. 2). This process enabled the rapid and user-friendly FE model generation and extraction of results. The details of the measurements and scaling of the template are explained in the supplementary material.

#### 2.3.2. Loading, boundary conditions, and finite element analyses

The outputs of the MS models used as inputs to the FE models consisted of [17, 23]: 1) knee flexion angle, 2) abduction/adduction and internal/external moments around the tibiofemoral joint (inverse dynamics), 3) abduction/adduction and internal/external moments generated by the muscles around the tibiofemoral joint, 4) flexion/extension, abduction/adduction, and internal/external moments generated by the quadriceps muscles around the patellofemoral joint, 5) tibiofemoral JCFs, and 6) patellofemoral JCFs (Fig. 1).

Tibiofemoral and patellofemoral JCFs (including the muscle forces) were applied to the femoral and patellar reference points (i.e., tibiofemoral and patellofemoral joints’ center of rotation), respectively (Fig. 1). Then the extracted muscle moment arms (explained in section 2.2.1) were used to calculate the moments generated by each muscle around the FE model reference points (i.e., around the joints’ center of rotation). The moments generated by each muscle were calculated by multiplying the muscle force by its moment arms separately for flexion-extension, abduction/adduction, and internal/external DoFs and for each time point of the trial. The calculated moments were then applied to the corresponding DoF of the reference point along with the knee external moments calculated from inverse dynamics (Fig. 1 and section 2.2.1).

The bottom of the tibia was fixed in all the FE models. All the nodes located on the femoral-cartilage to the subchondral-bone interface were coupled to the femoral reference point. Similarly, all the nodes located on the patellar-cartilage and subchondral-bone interface were coupled to the patellar reference point (Fig. 1). The knee flexion angle, knee joint moments (i.e., abduction/adduction and internal/external moments), and tibiofemoral JCFs were applied to the femoral reference point, and the patellar JCF and the moment generated by the quadriceps muscles (i.e., flexion/extension, abduction/adduction and internal/external moments) were applied to the patellar reference point (Fig. 1). The femur had 5 active DoFs, and the patella had 6 active DoFs in the FE models. More explanations about the loading and boundary conditions and inputs to the FE models are provided in the supplementary material (section 1.3.3, section 1.4, and Figs. S8 to S14) for both the EMG-assisted and SO-based MS-FE pipelines.

The aforementioned inputs to the FE models were automatically written to a file (one file per trial) and attached to the appropriate Abaqus input file (section 2.3.1) at the run time (detailed steps are explained in supplementary materials, section 1.4). Finally, the whole cycle of each trial/task was analyzed using Abaqus/Standard soils consolidation solver on an Intel(R) Xeon(R) CPU E5-2690 v3 (2.60GHz), single-core analysis.

### 2.4. Post-processing of the results and statistical analyses

The contact area, center of pressure (CoP), and average and maximum tissue mechanical responses, including maximum principal stress, collagen fibril strain, fluid pressure, and deviatoric and maximum shear strain were investigated within the femoral, tibial, and patellar cartilages and menisci. To calculate the average of tissue mechanical responses, for instance within the tibial surface, first all the nodes/elements of the tibial cartilage in contact with either femoral cartilage or menisci were selected separately at each time point of the cycles. Then the sum of nodal/elemental values of the parameter of interest was calculated and divided by the number of nodes/elements in the contact area for that time increment. The CoP on the tibial cartilage along an arbitrary axis (e.g., x) was calculated as follows:

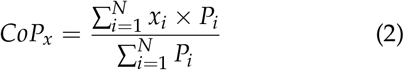

where *x_i_* is the coordinate of the center of the *i^th^* element in (arbitrary) *x* axis, *P_i_* is the average contact pressure within the *i^th^* element of the surface, and *N* is the total number of elements on the surface of interest.

The estimated results by the SO-based MS-FE models and the EMG-assisted MS-FE models were compared point-by-point (as a function of time) using statistical parametric mapping (SPM) paired t-tests [72], with p<0.05 and Bonferroni correction. Also, root mean square error (RMSE) and coefficient of determination (R^2^) between experimental and predicted muscle excitations were calculated for each MS modeling approach and separately for each subject’s trial (including all the time points).

## 3. Results

Using the developed pipeline, loading the MRIs (in MATLAB) and then measuring the morphological dimensions of each subject took only several minutes, from which the FRPVE FE model of each subject was created in several seconds. Executing the MS-FE analysis and delivering the results, on average, took 20 hours per one second for each activity (on a typical CPU and single-core analysis).

### 3.1. Muscle activations, joint kinematics, and joint kinetics

The estimated EMG-assisted muscle activations had fewer deviations from EMG envelopes compared to SO-based estimated muscle activations (Figs. S1 to S7). In 55% of all the activities, R2 and RMSE (expermental vs. predicted muscle activations) were significantly (p<0.05) different between the EMG-assisted and SO-based MS models. In 84% of these cases, t EMG-assisted MS model had significantly (p<0.05 higher R2 compared to the SO-based MS model (Figs 3 and S15).

**Fig. 3:**
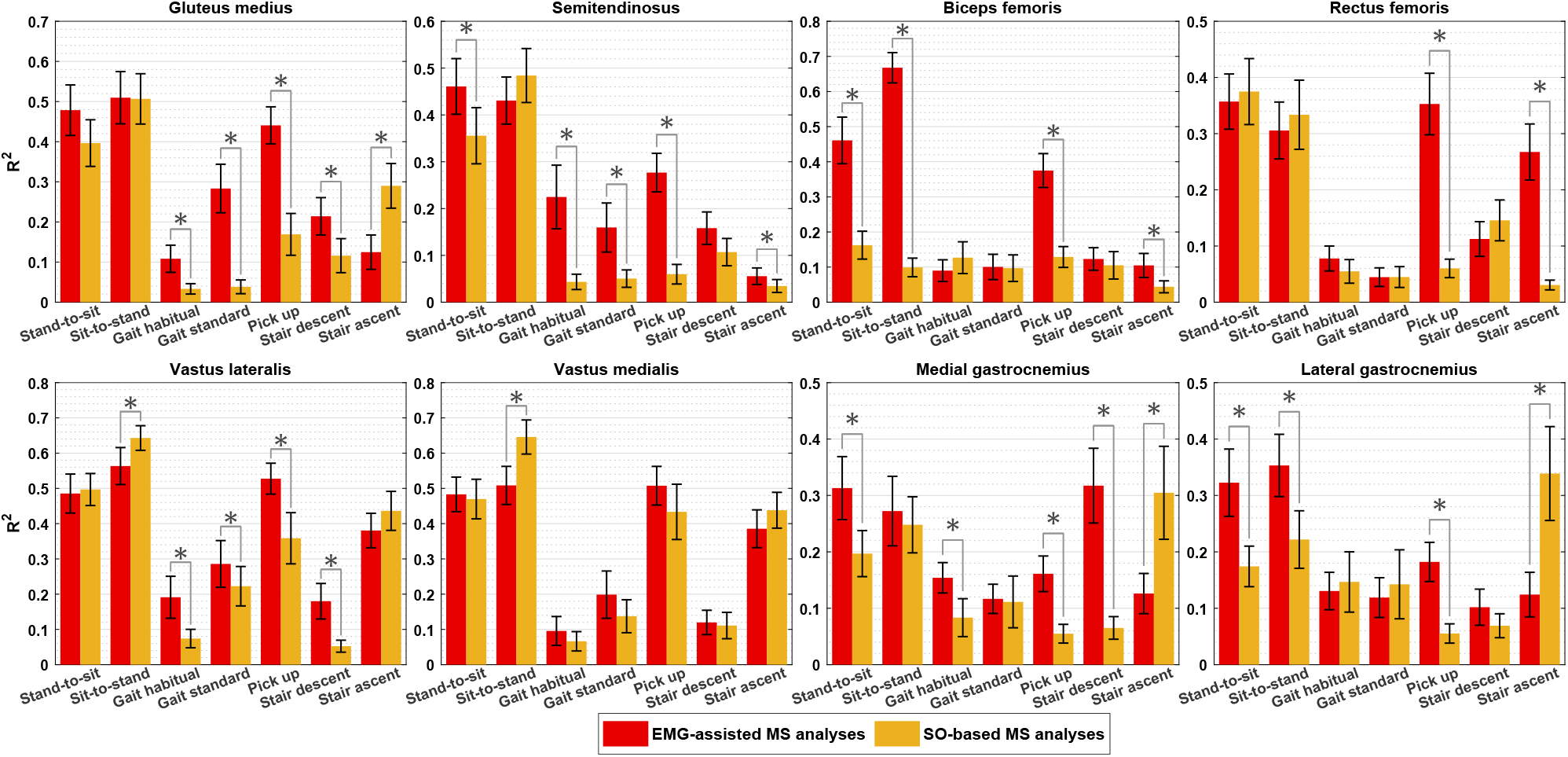
The coefficient of determination (R^2^) between the enveloped EMGs and estimated muscle activations by the EMG-assisted MS model (in red) and SO-based MS model (in yellow). Stars indicated significant differences (p<0.05) using paired sample t-test, and error bars show 95% confidence intervals.

The knee flexion angle, and abduction/adduction and internal/external rotation moments, from the EMG-assisted and SO-bas MS models were not significantly different (p<0.05 (Figs. S8 to S14). Nonetheless, the tibiofemoral JCFs including their peaks, in gait and stair negotiation estimated by the EMG-assisted MS model were significantly (p<0.05) higher than those of SO-based MS models for more than 70% of the cycles (Figs. S8 to S14). Further, the normalized JCF peaks of daily activities estimated by the EMG-assisted MS-FE pipeline bore a closer resemblance to the *in vivo* measured JCFs [15] compared to those of the SO-based MS-FE pipeline and those estimated by a SO-based 12 DoFs knee MS model reported previously [28] (Fig. 4). The EMG-assisted MS model estimated higher JCFs on the medial tibia than the lateral tibia (Fig. 5-A and B) during all the activities, although, the SO-based neural solution estimated higher JCF on the lateral tibia than medial tibia during stand-to-sit, sit-to-stand, and pick up (Fig. 5-C and D).

**Fig. 4:**
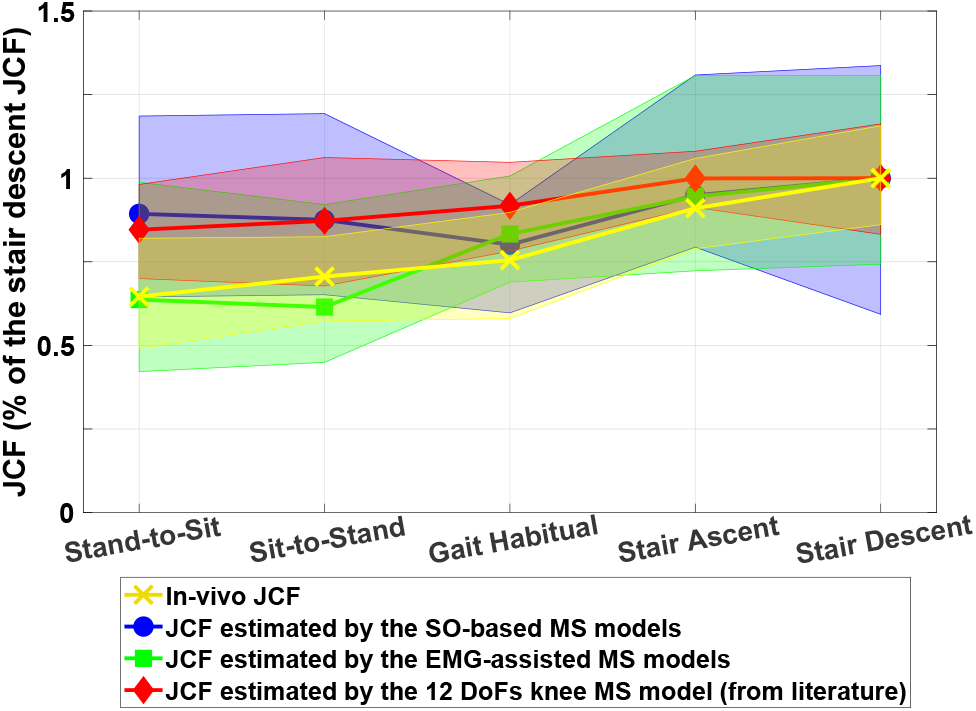
The peak JCF estimated by the SO-based and EMG-assisted MS models of the study compared to *in vivo* JCF [15] and those from the 12 DoFs knee MS model [28]. Note that the JCFs are normalized against the average stair descent JCF of each dataset, correspondingly.

**Fig. 5:**
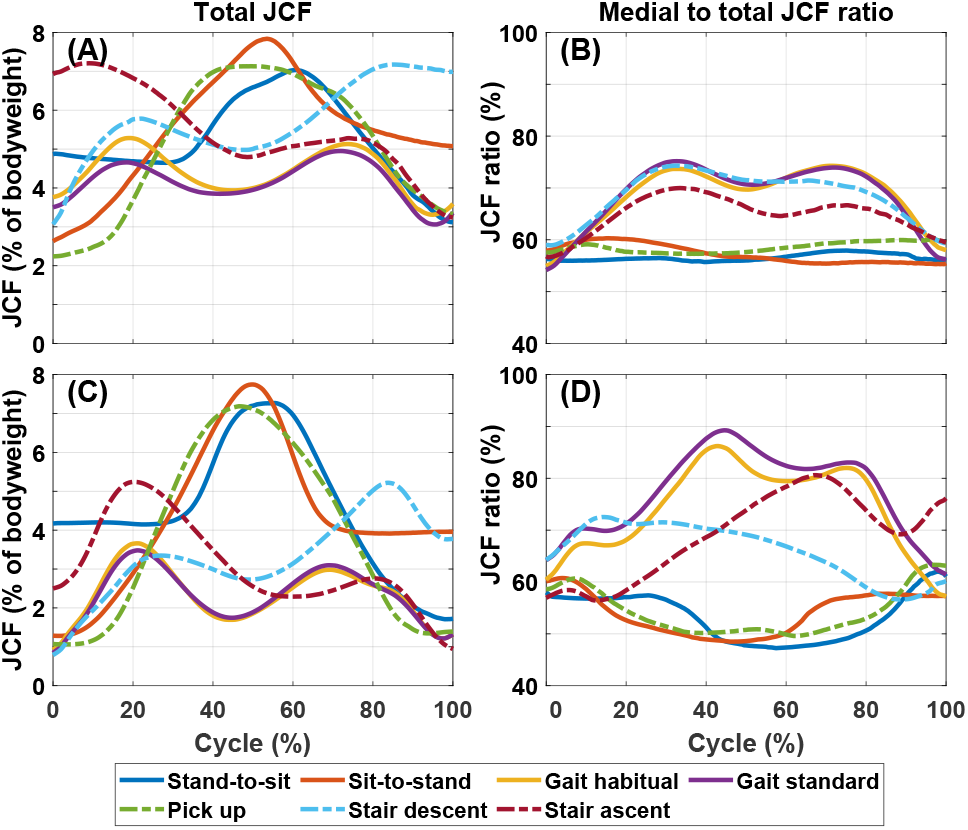
The total (A) and the medial-to-total ratio (B) of the tibiofemoral JCFs estimated by the EMG-assisted MS-FE models, and the total (C) and the medial-to-total ratio (D) of the tibiofemoral JCFs estimated by the SO-based MS-FE models. Plots report the 15 subject average profile for each activity. Deviations from the average (e.g., standard deviations) are not shown to improve the readability.

### 3.2. Contact pressure, contact area, and tissue mechanical responses

In general, more subject-specific variations were observed in the CoP at the maximum JCF among the subjects during the gait and pick up, compared to other daily activities (Fig. 6). Nonetheless, the CoP at the maximum JCF was consistent with those reported in previous *in situ* experiments and simulation-based studies reported for the gait and stair negotiation [14, 17, 73, 74].

**Fig. 6:**
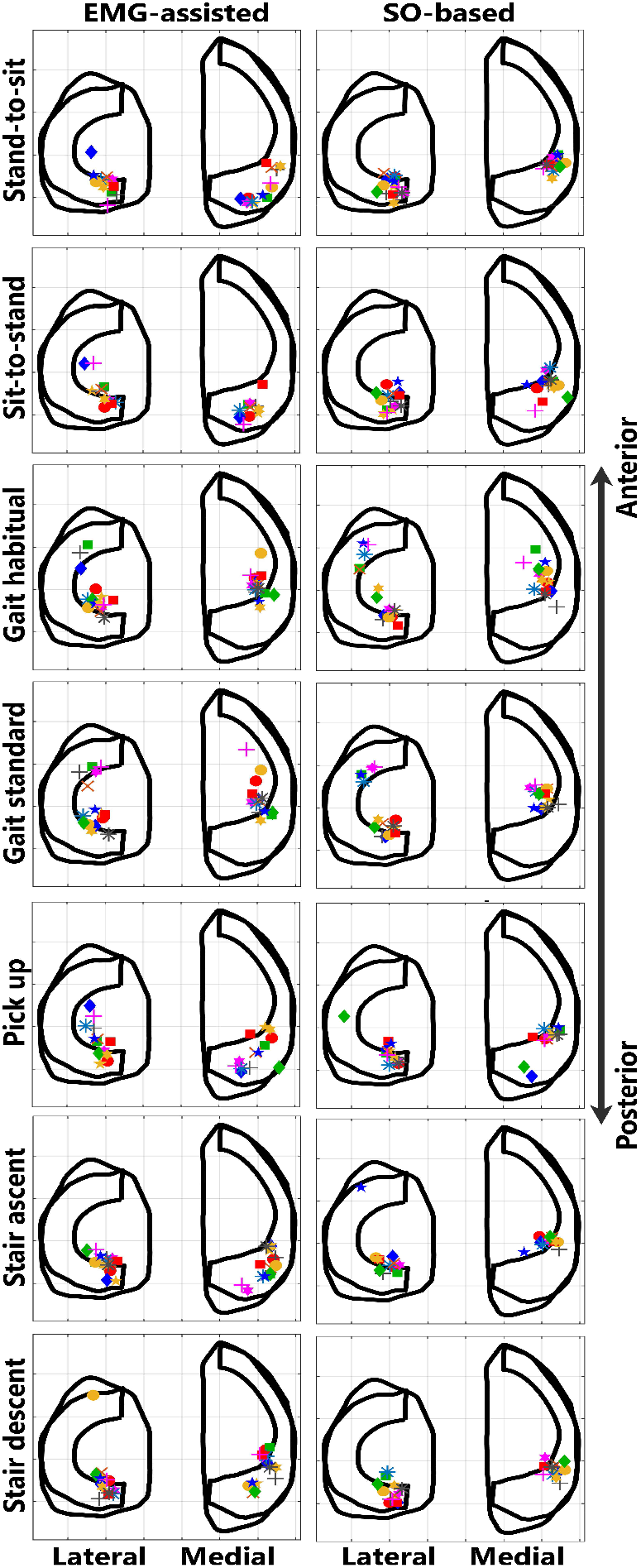
The tibial center of pressure (CoP) at the maximum JCF for 15 subjects of the study during the daily activities estimated by the EMG-assisted (on the left) and SO-based (on the right) MS-FE pipelines. Markers are representative of each subject of the study.

The medial and lateral tibial contact area during walking was significantly (p<0.05) different only for ~30% of the cycle between the EMG-assisted and the SO-based MS-FE models. In stair ascent, the contact area estimated by the EMG-assisted MS-FE model was significantly different (p<0.05) than that of the SO-based MS-FE model for ~20% and ~80% on the medial and lateral tibia, respectively. Nonetheless, there were fewer discrepancies between the contact area estimated by the EMG-assisted MS-FE model and those from *in situ* experiments [14] during walking and stair ascent (Fig. 7), compared to the SO-based MS-FE model.

**Fig. 7:**
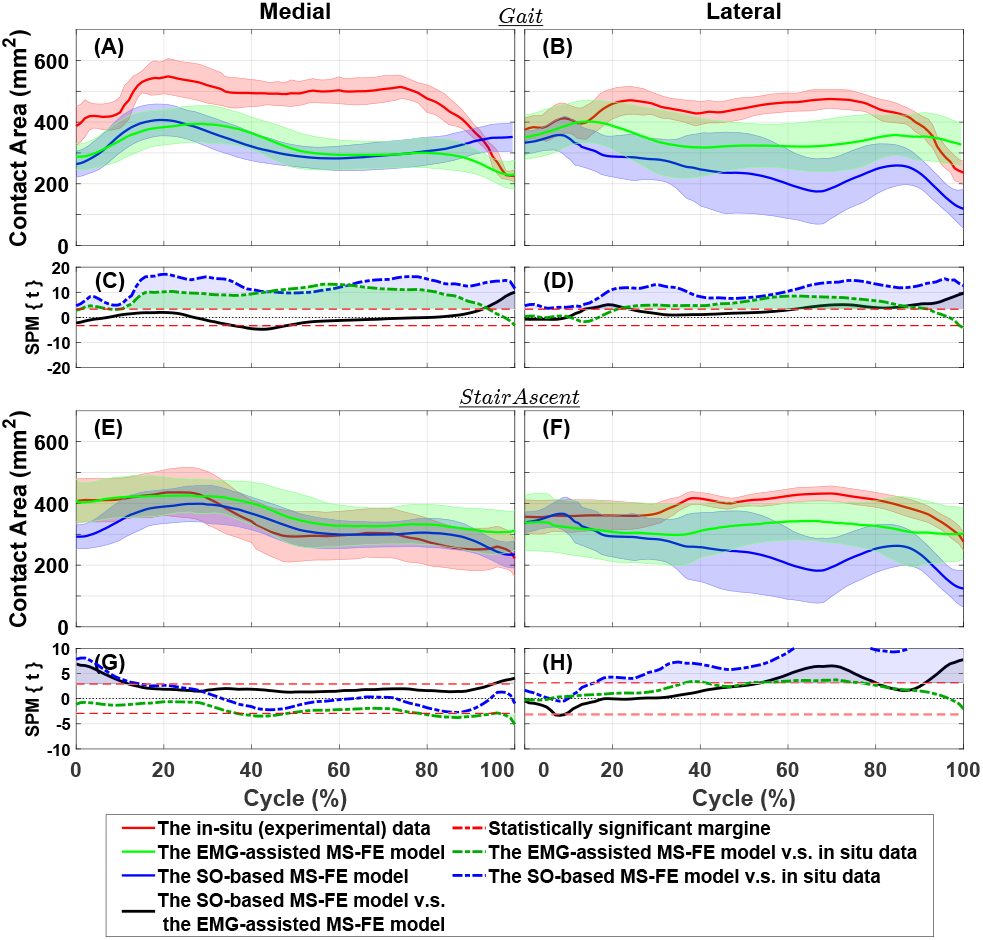
Contact area estimated by the EMG-assisted a SO-based MS-FE models of the study compared to experiments [14] for medial tibial cartilage during the gait (A) and stair ascent (E) and for lateral tibial cartilage during the gait (B) and stair ascent (F). For ease of comparison, simulation results are compared against each other (i.e., SO-based vs. EMG-assisted) using paired sample t-test and also against the experiments using independent samples t-test for medial tibial cartilage during the gait (C) and stair descent (G) and for lateral tibial cartilage during the gait (D) and stair ascent (H) using statistical parametric mapping.

The magnitudes and the mediolateral distribution of the estimated mechanical responses of cartilage during the gait were comparable with those reported in previous studies [17, 23, 29, 74, 75] (Figs. 8, S16 to S18). Within the lateral tibial cartilage, the average tissue mechanical responses were highest in stand-to-sit, sit-to-stand, and pick up and were lowest during walking in both the EMG-assisted and SO-based neural solutions (Figs. 8, S16, and S17, B and D). However, the maximum of the tissue mechanical responses was not substantially different between the activities within the medial tibial cartilage (Figs. 8, S16, and S17, A and C).

**Fig. 8:**
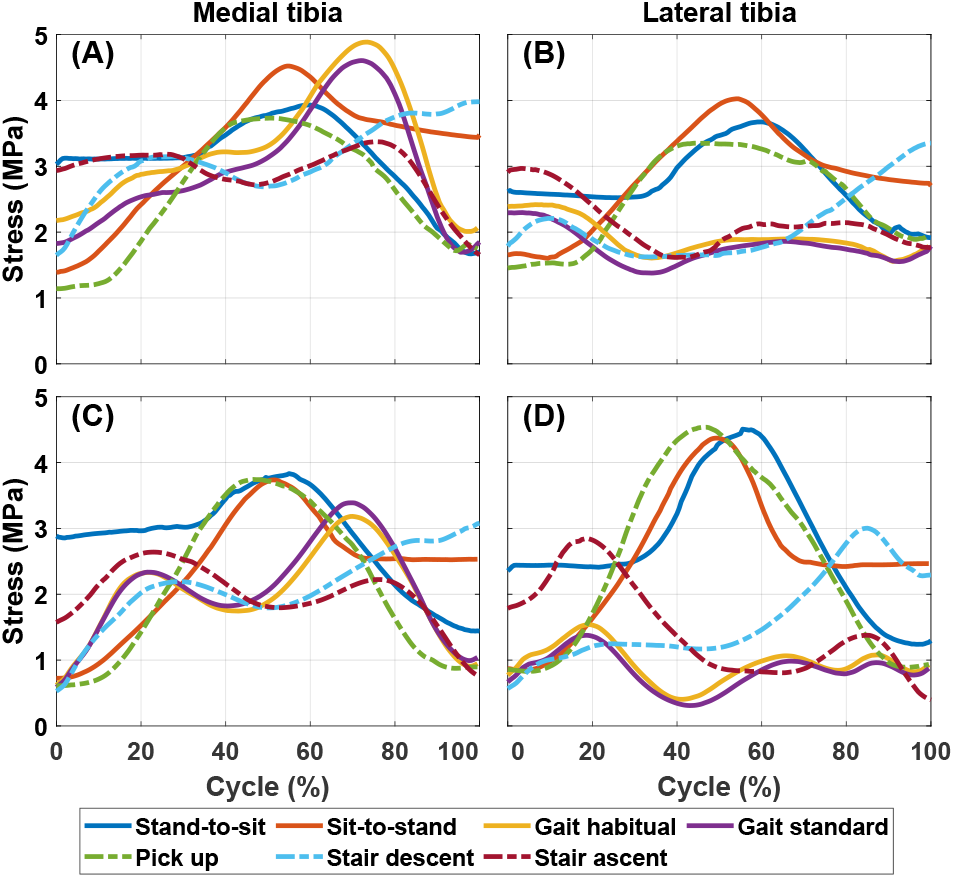
Average of the maximum principal stress estimated by the EMG-assisted MS-FE models on the medial tibia (A) and lateral tibia (B), and the SO-based MS-FE model on the medial tibia (C) and lateral tibia (D), reporting the 15 subject average profile for each activity. Deviations from the average (e.g., standard deviations) are not shown to improve the readability.

## 4. Discussion

### 4.1. Summary

In this study, a novel MS-FE modeling pipeline was established with a focus on feasibility for a rapid and user-friendly clinical assessment tool. Herein, a state-of-the-art EMG-assisted muscle-force driven FE model with FRP(V)E cartilages and menisci [17,23,49] was utilized. The EMG-assisted MS model enables the inclusion of subject-specific muscle activation patterns that may alter in subjects with MS disorders [24–27, 76, 77]. In addition, the FE model utilized a highly-detailed FRPVE soft tissue material model that has shown potentials if tissue-level mechanical responses, such as collagen fibril and nonfibrillar matrix strain, related to mechanically-induced cartilage adaptation and regeneration, are of interest [6, 37–41, 43, 46–48]. To assess and validate the developed pipeline, we investigated the knee joint loading and tissue mechanical responses during different daily activities of the individuals with KOA. Besides, the results of the EMG-assisted MS-FE models were compared to those of the SO-based models to investigate potential alterations in the analyzed parameters due to different MS analysis approaches.

### 4.2. The atlas-based MS-FE modeling toolbox

For the first time, we introduced and validated a semi-automated and rapid MS-FE analysis toolbox capable of modeling the whole knee joint, incorporating subject-specific muscle activation patterns, joint kinematics and kinetics, and multiscale tissue mechanics. The presented pipeline took less than a day to create the models, perform analyses of a general task, and then deliver the results. Except for scaling the MS and FE models, the rest of the pipeline, including running MS analyses in both OpenSim and CEINMS, writing inputs to the FE models, executing the FE models, and then extracting the results, does not require user interactions. Of course, possible convergence difficulties in the FE model and interpretation and verification of the results of each step still require supervision and expertise, although this could be automated in the future. A similar workflow without the atlas-based modeling toolbox requires a high level of unique skills with several months of training to perform segmentation and meshing, incorporate the FRPVE materials model, interconnect the models, and get models to converge. Even for an expert-user, those are laborious tasks taking several weeks/months to perform manually [78]. Utilizing image processing and machine learning techniques [79], our further studies aim to reduce the simulation time even towards real-time EMG-assisted MS-FE analyses (e.g., as done for the Achilles tendon, [80]) and to make the whole pipeline automatic (i.e., scaling and morphing of the MS and FE models’ geometries) [81].

### 4.3. SO-based compared to EMG-assisted MS-FE analyses

In our previous study [17], the EMG-assisted MS-FE modeling workflow bore a closer resemblance to the experiments, compared to the SO-based MS-FE model. As a further evaluation, we compared the results obtained from the SO-based and EMG-assisted MS-FE analyses using the dataset of the current study. The enveloped EMGs showed a wide variation between subjects (Figs. S1 to S7) that may account for individual variations in muscle recruitment strategies [24–27,77]. These variations were also seen in the muscle activations estimated by the EMG-assisted MS model but to a lesser degree in those estimated by the SO-based MS model (Figs. S1 to S7). Consequently, more variations were observed in estimated JCFs and tissue mechanical responses using the EMG-assisted MS-FE model compared to the SO-based model (Figs. 4, and S8 to S14).

Higher knee adduction moment during stand-to-sit, sit-to-stand, and pick up compared to other activities (Figs. S8 to S14) accounts for the lowest medial-to-total JCF ratio at the peak of the total JCF in these three activities compared to other daily activities (Fig. 5-B and D), both the EMG-assisted and SO-based MS models). More importantly, the SO-based MS model estimated higher JCF peaks on the lateral tibia than the medial tibia during stand-to-sit, sit-to-stand, and pick up, as opposed to the EMG-assisted MS model. This may be due to significantly (p<0.05) higher activation of the biceps femoris in the SO-based MS models in contrast with those of the EMG-assisted models and the measured EMGs (Figs. 3, S1, S2, and S5).

Previous studies [24–27, 76, 77] reported higher (up to double) muscle activations and co-contractions in subjects with KOA compared to healthy individuals. Hence, assisting the MS analyses with EMGs and considering higher correlations between the EMG envelopes and the muscle activations estimated by the EMG-assisted MS models than those of the SO-based models (Figs. 3 and S1 to S7) explain higher JCFs estimated by the EMG-assisted MS models of the study, compared to those estimated by the SO-based MS models (Figs. 5 and S8 to S14). Interestingly, the maximum JCF of the daily activities estimated by the EMG-assisted MS model were more consistent with those from experiments compared to the SO-based estimated JCFs with either 1 DoF or 12 DoFs knee models (Fig. 4). In line with our results, it has been shown [82] that the ability of SO-based MS model with a 1 DoF knee to predict the knee JCFs in different subjects and activities (other than gait) is limited, compared to EMGs and *in vivo* JCFs.

To summarize, the above-mentioned variations and differences, consistent with previous studies [24–27, 76, 77, 82, 83], may emphasize the necessity of assisting the MS model with subject-specific muscle activation patterns, especially in individuals with MS disorders and pain.

### 4.4. Limitations

We did not group participants according to KOA grade, pain score, etc. Moreover, this study can be seen as the initial assessment of the developed MS-FE analysis pipeline. However, our results showed that the developed workflow has the potential to analyze multi-level joint mechanical responses of subjects with different, e.g., KOA grade due to the inclusion of subject-specific kinematics, kinetics, muscle activation patterns, and joint geometries. Yet, complimentary evaluations with larger cohorts and more activities are needed to further evaluate the developed pipeline.

The muscle-tendon parameters of the utilized MS models were not subject-specific. Nonetheless, the calibration module of the CEINMS using the subject’s EMG envelopes is shown to attenuate the effect of the muscle-tendon uncertainties on the, e.g., estimated JCFs [83]. Also, material parameters of the knee soft tissues, including those of cartilages, may vary at different sites [84, 85] that can alter the magnitude of the local tissue mechanical responses [86, 87]. However, no practical methods are available to fully extract subject-specific mechanical properties of knee soft tissues, and hence, the soft tissue material parameters utilized in this study (Table S2) were adopted from the literature. Nonetheless, the FE model of the study has shown the potentials to incorporate subject- and site-specific material parameters [88–90].

The use of one template (compared to, e.g., choosing among several templates) is suggested before [49]; however, our future studies aim to enhance the atlas-based FE modeling method employing multi-template approaches and nonlinear scaling methods such as statistical shape modeling. The atlas-based FE modeling approach has been favorably evaluated and validated against the follow-up data (i.e., KOA progression) [49], although it may be seen as a limitation of this approach, especially in the subjects with evident cartilage lesions. Nevertheless, the joint-space narrowing and the cartilage thickness were reflected in the morphological measurements of each subject. Though, estimation of the local tissue mechanical responses around the cartilage lesion, in general, may be addressed as a limitation of the atlas-based modeling approach. Currently, the inclusion of subject-specific cartilage lesions requires lengthy manual mesh editing.

### 4.5. Applications and further developments

This study showcases the usability of the developed pipeline for various tasks and patients in different levels of KOA, as well as the potential for implementing the pipeline as a feasible and rapid MS-FE analysis toolbox not only for research purposes with large cohorts, but also for clinical applications.

The stress and strain within the fibrillar (collagen) and nonfibrillar (proteoglycans) matrices of the cartilage and menisci are reported as indicators governing the cartilage adaptation and degradation response [43, 46, 47]. Hence, our workflow employing the FRPVE material model may promisingly be used in the subject-specific modification of different activities and the design of rehabilitation protocols, to slow the onset or progression of the KOA according to the estimated subject-specific joint mechanics. Also, our further study aims to investigate the knee joint mechanics of KOA individuals in different daily activities and rehabilitation exercises in more detail and explores the correlation between the subject’s regional pain intensity and localized joint mechanics.

This study showcases the usability of the developed pipeline for various tasks and patients in different levels of KOA, as well as the potential for implementing the pipeline as a feasible and rapid MS-FE analysis toolbox not only for research purposes with large cohorts, but also for clinical applications. A few models have been developed to predict cartilage degeneration and KOA progression based on the mechanobiological response of the joint’s soft tissue [6, 9, 46, 47, 91]. For instance, collagen fibril strain (related to, e.g., collagen damage) and fluid flow or maximum shear strain (related to, e.g., cell death and fixed charged density loss of the proteoglycans within the cartilage) are reported as measures for the prediction of cartilage adaptation and degradation response, consistent with experiments [6, 9, 46, 47, 91]. Nonetheless, none of these studies [6,9,46,47,91] have been incorporated into a multiscale MS-FE model considering subject-specific joint loading. Integrating the EMG-assisted MS-FE pipeline of our study with cartilage remodeling algorithms [6, 9, 46, 47, 91], as a part of our further research, may bring more accuracy when subject-specific mechanically-induced soft tissue adaptation and degeneration is of interest. As a result, the developed workflow can potentially be used in the subject-specific modification of different activities and the design of rehabilitation protocols to slow the onset or progression of the KOA according to the estimated subject’s joint mechanics.

### 4.6. Conclusion

For the first time, a semi-automated rapid MS-FE analysis pipeline was developed and validated against experimental data. The muscle activation patterns and knee joint mechanics in different size scales were estimated and reported by EMG-assisted and SO-based MS-FE modeling of common daily activities in individuals with KOA. This offers a comprehensive source of locomotion information. In general, the EMG-assisted MS-FE pipeline bore a closer resemblance to experiments, compared to the SO-based MS-FE pipeline. Hence, our results emphasize the importance of assisting the MS-FE analysis with subjects’ muscle activation patterns, especially when simulating different physical activities of KOA subjects. More importantly, the developed pipeline showed great potentials as a rapid MS-FE analysis toolbox to investigate multiscale knee mechanics during different activities of individuals with KOA. Nonetheless, our future research aims to investigate the feasibility of the pipeline to personalize daily activity routines and rehabilitation protocols of KOA individuals, e.g., to slow the KOA progress by optimal loading of knee load-bearing tissue.

## Acknowledgment

This study was financially supported by the European Union’s Horizon 2020 research and innovation program under the Marie Sklodowska-Curie grant agreement No. 713645, Academy of Finland (grant no. 324529, 324994, 328920), Sigrid Juselius Foundation and Business Finland (grant no. 3455/31/2019), the Finnish Cultural Foundation and the Päivikki and Sakari Sohlberg Foundation, the European Regional Developments Fund and the University of Eastern Finland under the project: Human measurement and analysis - research and innovation laboratories (HUMEA, project identifiers: A73200 and A73241). This study was also partly funded by a grant from the National Health and Medical Research Council of Australia (2001734) to DGL and RKK.

## Notes

### Competing Interest Statement

The authors have declared no competing interest.

## References

[1] S. Van Grinsven, R. Van Cingel, C. Holla, and C. Van Loon, “Evidence-based rehabilitation following anterior cruciate ligament reconstruction,” Knee Surgery, Sports Traumatology, Arthroscopy, vol. 18, no. 8, pp. 1128–1144, 2010.

[2] E. Alentorn-Geli, G. D. Myer, H. J. Silvers, G. Samitier, D. Romero, C. Lázaro-Haro, and R. Cugat, “Prevention of non-contact anterior cruciate ligament injuries in soccer players. part 1: Mechanisms of injury and underlying risk factors,” Knee surgery, sports traumatology, arthroscopy, vol. 17, no. 7, pp. 705–729, 2009.

[3] A. Baliunas, D. Hurwitz, A. Ryals, A. Karrar, J. Case, J. Block, and T. Andriacchi, “Increased knee joint loads during walking are present in subjects with knee osteoarthritis,” Osteoarthritis and cartilage, vol. 10, no. 7, pp. 573–579, 2002.

[4] C. Egloff, T. Hügle, and V. Valderrabano, “Biomechanics and pathomechanisms of osteoarthritis,” Swiss medical weekly, vol. 142, no. 2930, 2012.

[5] D. Felson, “Osteoarthritis as a disease of mechanics,” Osteoarthritis and Cartilage, vol. 21, no. 1, pp. 10–15, 2013.

[6] M. K. Liukkonen, M. E. Mononen, P. Vartiainen, P. Kaukinen, T. Bragge, J.-S. Suomalainen, M. K. Malo, S. Venesmaa, P. Käkelä, J. Pihlajamäki et al., “Evaluation of the effect of bariatric surgery-induced weight loss on knee gait and cartilage degeneration,” Journal of biomechanical engineering, vol. 140, no. 4, 2018.

[7] T. Miyazaki, M. Wada, H. Kawahara, M. Sato, H. Baba, and S. Shimada, “Dynamic load at baseline can predict radiographic disease progression in medial compartment knee osteoarthritis,” Annals of the rheumatic diseases, vol. 61, no. 7, pp. 617–622, 2002.

[8] B. T. Raines, E. Naclerio, and S. L. Sherman, “Management of anterior cruciate ligament injury: what’s in and what’s out?” Indian journal of orthopaedics, vol. 51, pp. 563–575, 2017.

[9] W. Wilson, C. van Burken, C. van Donkelaar, P. Buma, B. van Rietbergen, and R. Huiskes, “Causes of mechanically induced collagen damage in articular cartilage,” Journal of Orthopaedic Research, vol. 24, no. 2, pp. 220–228, 2006.

[10] J. E. Bischof, C. E. Spritzer, A. M. Caputo, M. E. Easley, J. K. DeOrio, J. A. Nunley II, and L. E. DeFrate, “In vivo cartilage contact strains in patients with lateral ankle instability,” Journal of biomechanics, vol. 43, no. 13, pp. 2561–2566, 2010.

[11] K. Connolly, J. Ronsky, L. Westover, J. Küpper, and R. Frayne, “Differences in patellofemoral contact mechanics associated with patellofemoral pain syndrome,” Journal of biomechanics, vol. 42, no. 16, pp. 2802–2807, 2009.

[12] D. D. D’Lima, S. Patil, N. Steklov, J. E. Slamin, and C. W. Colwell Jr, “Tibial forces measured in vivo after total knee arthroplasty,” The Journal of arthroplasty, vol. 21, no. 2, pp. 255–262, 2006.

[13] B. J. Fregly, T. F. Besier, D. G. Lloyd, S. L. Delp, S. A. Banks, M. G. Pandy, and D. D. D’lima, “Grand challenge competition to predict in vivo knee loads,” Journal of orthopaedic research, vol. 30, no. 4, pp. 503–513, 2012.

[14] S. Gilbert, T. Chen, I. D. Hutchinson, D. Choi, C. Voigt, R. F. Warren, and S. A. Maher, “Dynamic contact mechanics on the tibial plateau of the human knee during activities of daily living,” Journal of biomechanics, vol. 47, no. 9, pp. 2006–2012, 2014.

[15] I. Kutzner, B. Heinlein, F. Graichen, A. Bender, A. Rohlmann, A. Halder, A. Beier, and G. Bergmann, “Loading of the knee joint during activities of daily living measured in vivo in five subjects,” Journal of biomechanics, vol. 43, no. 11, pp. 2164–2173, 2010.

[16] A. Mündermann, C. O. Dyrby, D. D. D’Lima, C. W. Colwell Jr, and T. P. Andriacchi, “In vivo knee loading characteristics during activities of daily living as measured by an instrumented total knee replacement,” Journal of Orthopaedic Research, vol. 26, no. 9, pp. 1167–1172, 2008.

[17] A. Esrafilian, L. Stenroth, M. E. Mononen, P. Tanska, S. Van Rossom, D. G. Lloyd, I. Jonkers, and R. K. Korhonen, “12 degrees of freedom muscle force driven fibril-reinforced poroviscoelastic finite element model of the knee joint,” IEEE Transactions on Neural Systems and Rehabilitation Engineering, vol. 29, pp. 123–133, 2021.

[18] R. L. Lenhart, J. Kaiser, C. R. Smith, and D. G. Thelen, “Prediction and validation of load-dependent behavior of the tibiofemoral and patellofemoral joints during movement,” Annals of biomedical engineering, vol. 43, no. 11, pp. 2675–2685, 2015.

[19] H. Marouane, A. Shirazi-Adl, and M. Adouni, “3d active-passive response of human knee joint in gait is markedly altered when simulated as a planar 2d joint,” Biomechanics and modeling in mechanobiology, vol. 16, no. 2, pp. 693–703, 2017.

[20] M. A. Marra, V. Vanheule, R. Fluit, B. H. Koopman, J. Rasmussen, N. Verdonschot, and M. S. Andersen, “A subject-specific musculoskeletal modeling framework to predict in vivo mechanics of total knee arthroplasty,” Journal of biomechanical engineering, vol. 137, no. 2, 2015.

[21] A. Navacchia, D. R. Hume, P. J. Rullkoetter, and K. B. Shelburne, “A computationally efficient strategy to estimate muscle forces in a finite element musculoskeletal model of the lower limb,” Journal of biomechanics, vol. 84, pp. 94–102, 2019.

[22] C. Pizzolato, D. G. Lloyd, M. Sartori, E. Ceseracciu, T. F. Besier, B. J. Fregly, and M. Reggiani, “Ceinms: A toolbox to investigate the influence of different neural control solutions on the prediction of muscle excitation and joint moments during dynamic motor tasks,” Journal of biomechanics, vol. 48, no. 14, pp. 3929–3936, 2015.

[23] A. Esrafilian, L. Stenroth, M. Mononen, P. Tanska, J. Avela, and R. Korhonen, “emg-assisted muscle force driven finite element model of the knee joint with fibril-reinforced poroelastic cartilages and menisci,” Scientific reports, vol. 10, no. 1, pp. 1–16, 2020.

[24] T. L. Heiden, D. G. Lloyd, and T. R. Ackland, “Knee joint kinematics, kinetics and muscle co-contraction in knee osteoarthritis patient gait,” Clinical biomechanics, vol. 24, no. 10, pp. 833–841, 2009.

[25] C. L. Hubley-Kozey, N. A. Hill, D. J. Rutherford, M. J. Dunbar, and W. D. Stanish, “Co-activation differences in lower limb muscles between asymptomatic controls and those with varying degrees of knee osteoarthritis during walking,” Clinical biomechanics, vol. 24, no. 5, pp. 407–414, 2009.

[26] B. Killen, D. Saxby, K. Fortin, B. Gardiner, T. Wrigley, A. Bryant, and D. Lloyd, “Individual muscle contributions to tibiofemoral compressive articular loading during walking, running and sidestepping,” Journal of biomechanics, vol. 80, pp. 23–31, 2018.

[27] L. C. Schmitt and K. S. Rudolph, “Muscle stabilization strategies in people with medial knee osteoarthritis: the effect of instability,” Journal of Orthopaedic Research, vol. 26, no. 9, pp. 1180–1185, 2008.

[28] S. Van Rossom, C. R. Smith, D. G. Thelen, B. Van-wanseele, D. Van Assche, and I. Jonkers, “Knee joint loading in healthy adults during functional exercises: implications for rehabilitation guidelines,” journal of orthopaedic & sports physical therapy, vol. 48, no. 3, pp. 162–173, 2018.

[29] K. Halonen, C. M. Dzialo, M. Mannisi, M. Venäläinen, M. de Zee, and M. S. Andersen, “Workflow assessing the effect of gait alterations on stresses in the medial tibial cartilage-combined musculoskeletal modelling and finite element analysis,” Scientific reports, vol. 7, no. 1, pp. 1–14, 2017.

[30] C. Pizzolato, M. Reggiani, D. J. Saxby, E. Ceseracciu, L. Modenese, and D. G. Lloyd, “Biofeedback for gait retraining based on real-time estimation of tibiofemoral joint contact forces,” IEEE Transactions on Neural Systems and Rehabilitation Engineering, vol. 25, no. 9, pp. 1612–1621, 2017.

[31] C. M. Dzialo, M. Mannisi, K. Halonen, M. de Zee, J. Woodburn, and M. S. Andersen, “Gait alteration strategies for knee osteoarthritis: a comparison of joint loading via generic and patient-specific musculoskeletal model scaling techniques,” International Biomechanics, vol. 6, no. 1, pp. 54–65, 2019.

[32] S. C. Starkey, G. K. Lenton, D. J. Saxby, R. S. Hinman, K. L. Bennell, T. Wrigley, D. Lloyd, and M. Hall, “Effect of exercise on knee joint contact forces in people following medial partial meniscectomy: A secondary analysis of a randomised controlled trial,” Gait & Posture, vol. 79, pp. 203–209, 2020.

[33] P. Gerus, M. Sartori, T. F. Besier, B. J. Fregly, S. L. Delp, S. A. Banks, M. G. Pandy, D. D. D’Lima, and D. G. Lloyd, “Subject-specific knee joint geometry improves predictions of medial tibiofemoral contact forces,” Journal of biomechanics, vol. 46, no. 16, pp. 2778–2786, 2013.

[34] J. Ihn, S. Kim, and I. Park, “In vitro study of contact area and pressure distribution in the human knee after partial and total meniscectomy,” International orthopaedics, vol. 17, no. 4, pp. 214–218, 1993.

[35] M. Kelly, D. Fithian, K. Chern, and V. C. Mow, “Structure and function of the meniscus: basic and clinical implications,” in Biomechanics of diarthrodial joints. Springer, 1990, pp. 191–211.

[36] E. L. Radin, F. de Lamotte, and P. Maquet, “Role of the menisci in the distribution of stress in the knee.” Clinical orthopaedics and related research, no. 185, pp. 290–294, 1984.

[37] O. Klets, M. E. Mononen, P. Tanska, M. T. Nieminen, R. K. Korhonen, and S. Saarakkala, “Comparison of different material models of articular cartilage in 3d computational modeling of the knee: Data from the osteoarthritis initiative (oai),” Journal of biomechanics, vol. 49, no. 16, pp. 3891–3900, 2016.

[38] L. Li and K. Gu, “Reconsideration on the use of elastic models to predict the instantaneous load response of the knee joint,” Proceedings of the Institution of Mechanical Engineers, Part H: Journal of Engineering in Medicine, vol. 225, no. 9, pp. 888–896, 2011.

[39] J. Mäkelä, S. Han, W. Herzog, and R. Korhonen, “Very early osteoarthritis changes sensitively fluid flow properties of articular cartilage,” Journal of biomechanics, vol. 48, no. 12, pp. 3369–3376, 2015.

[40] J. P. Quiroga, W. Wilson, K. Ito, and C. Van Donkelaar, “Relative contribution of articular cartilage’s constitutive components to load support depending on strain rate,” Biomechanics and modeling in mechanobiology, vol. 16, no. 1, pp. 151–158, 2017.

[41] A. Shirazi-Adl, “On the fibre composite material models of disc annulus—comparison of predicted stresses,” Journal of Biomechanics, vol. 22, no. 4, pp. 357–365, 1989.

[42] G. A. Orozco, P. Tanska, C. Florea, A. J. Grodzinsky, and R. K. Korhonen, “A novel mechanobiological model can predict how physiologically relevant dynamic loading causes proteoglycan loss in mechanically injured articular cartilage,” Scientific reports, vol. 8, no. 1, pp. 1–16, 2018.

[43] S. Hosseini, W. Wilson, K. Ito, and C. Van Donkelaar, “A numerical model to study mechanically induced initiation and progression of damage in articular cartilage,” Osteoarthritis and cartilage, vol. 22, no. 1, pp. 95–103, 2014.

[44] A. Chen, W. Bae, R. Schinagl, and R. Sah, “Depth- and strain-dependent mechanical and electromechanical properties of full-thickness bovine articular cartilage in confined compression,” Journal of biomechanics, vol. 34, no. 1, pp. 1–12, 2001.

[45] N. Mukherjee and J. Wayne, “Load sharing between solid and fluid phases in articular cartilage: Ii—comparison of experimental results and up finite element predictions,” 1998.

[46] G. A. Orozco, P. Bolcos, A. Mohammadi, M. S. Tanaka, M. Yang, T. M. Link, B. Ma, X. Li, P. Tanska, and R. K. Korhonen, “Prediction of local fixed charge density loss in cartilage following acl injury and reconstruction: A computational proof-of-concept study with mri follow-up,” Journal of Orthopaedic Research®, 2020.

[47] A. S. Eskelinen, P. Tanska, C. Florea, G. A. Orozco, P. Julkunen, A. J. Grodzinsky, and R. K. Korhonen, “Mechanobiological model for simulation of injured cartilage degradation via pro-inflammatory cytokines and mechanical stimulus,” PLoS computational biology, vol. 16, no. 6, p. e1007998, 2020.

[48] P. O. Bolcos, M. E. Mononen, M. S. Tanaka, M. Yang, J.-S. Suomalainen, M. J. Nissi, J. Töyräs, B. Ma, X. Li, and R. K. Korhonen, “Identification of locations susceptible to osteoarthritis in patients with anterior cruciate ligament reconstruction: Combining knee joint computational modelling with follow-up t1*ρ* and t2 imaging,” Clinical Biomechanics, vol. 79, p. 104844, 2020.

[49] M. E. Mononen, M. K. Liukkonen, and R. K. Korhonen, “Utilizing atlas-based modeling to predict knee joint cartilage degeneration: data from the osteoarthritis initiative,” Annals of biomedical engineering, vol. 47, no. 3, pp. 813–825, 2019.

[50] R. Altman, E. Asch, D. Bloch, G. Bole, D. Borenstein, K. Brandt, W. Christy, T. Cooke, R. Greenwald, M. Hochberg et al., “Development of criteria for the classification and reporting of osteoarthritis: classification of osteoarthritis of the knee,” Arthritis & Rheumatism: Official Journal of the American College of Rheumatology, vol. 29, no. 8, pp. 1039–1049, 1986.

[51] H. J. Hermens, B. Freriks, R. Merletti, D. Stegeman, J. Blok, G. Rau, C. Disselhorst-Klug, and G. Hägg, “European recommendations for surface electromyography,” Roessingh research and development, vol. 8, no. 2, pp. 13–54, 1999.

[52] A. Mantoan, C. Pizzolato, M. Sartori, Z. Sawacha, C. Cobelli, and M. Reggiani, “Motonms: A matlab toolbox to process motion data for neuromusculoskeletal modeling and simulation,” Source code for biology and medicine, vol. 10, no. 1, pp. 1–14, 2015.

[53] D. G. Lloyd and T. F. Besier, “An emg-driven musculoskeletal model to estimate muscle forces and knee joint moments in vivo,” Journal of biomechanics, vol. 36, no. 6, pp. 765–776, 2003.

[54] D. S. Catelli, M. Wesseling, I. Jonkers, and M. Lamontagne, “A musculoskeletal model customized for squatting task,” Computer methods in biomechanics and biomedical engineering, vol. 22, no. 1, pp. 21–24, 2019.

[55] S. L. Delp, F. C. Anderson, A. S. Arnold, P. Loan, A. Habib, C. T. John, E. Guendelman, and D. G. Thelen, “Opensim: open-source software to create and analyze dynamic simulations of movement,” IEEE transactions on biomedical engineering, vol. 54, no. 11, pp. 1940–1950, 2007.

[56] M. Sartori, M. Reggiani, D. Farina, and D. G. Lloyd, “Emg-driven forward-dynamic estimation of muscle force and joint moment about multiple degrees of freedom in the human lower extremity,” PloS one, vol. 7, no. 12, p. e52618, 2012.

[57] T. S. Buchanan, D. G. Lloyd, K. Manal, and T. F. Besier, “Neuromusculoskeletal modeling: estimation of muscle forces and joint moments and movements from measurements of neural command,” Journal of applied biomechanics, vol. 20, no. 4, pp. 367–395, 2004.

[58] M. Sartori, D. Farina, and D. G. Lloyd, “Hybrid neuromusculoskeletal modeling to best track joint moments using a balance between muscle excitations derived from electromyograms and optimization,” Journal of biomechanics, vol. 47, no. 15, pp. 3613–3621, 2014.

[59] S. Below, S. P. Arnoczky, J. Dodds, C. Kooima, and N. Walter, “The split-line pattern of the distal femur: A consideration in the orientation of autologous cartilage grafts,” Arthroscopy: The Journal of Arthroscopic & Related Surgery, vol. 18, no. 6, pp. 613–617, 2002.

[60] A. Benninghoff, “Form und bau der gelenkknorpel in ihren beziehungen zur funktion,” Zeitschrift für Zellforschung und mikroskopische Anatomie, vol. 2, no. 5, pp. 783–862, 1925.

[61] P. Boettcher, M. Zeissler, J. Maierl, V. Grevel, and G. Oechtering, “Mapping of split-line pattern and cartilage thickness of selected donor and recipient sites for autologous osteochondral transplantation in the canine stifle joint,” Veterinary surgery, vol. 38, no. 6, pp. 696–704, 2009.

[62] D. W. Goodwin, Y. Z. Wadghiri, H. Zhu, C. J. Vinton, E. D. Smith, and J. F. Dunn, “Macroscopic structure of articular cartilage of the tibial plateau: influence of a characteristic matrix architecture on mri appearance,” American Journal of Roentgenology, vol. 182, no. 2, pp. 311–318, 2004.

[63] P. Julkunen, P. Kiviranta, W. Wilson, J. S. Jurvelin, and R. K. Korhonen, “Characterization of articular cartilage by combining microscopic analysis with a fibril-reinforced finite-element model,” Journal of biomechanics, vol. 40, no. 8, pp. 1862–1870, 2007.

[64] W. Wilson, C. van Donkelaar, B. van Rietbergen, K. Ito, and R. Huiskes, “Erratum to “stresses in the local collagen network of articular cartilage: a poroviscoelastic fibril-reinforced finite element study” [journal of biomechanics 37 (2004) 357–366] and “a fibril-reinforced poroviscoelastic swelling model for articular cartilage” [journal of biomechanics 38 (2005) 1195–1204],” Journal of Biomechanics, vol. 38, no. 10, pp. 2138–2140, 2005.

[65] W. Wilson, C. Van Donkelaar, B. Van Rietbergen, K. Ito, and R. Huiskes, “Stresses in the local collagen network of articular cartilage: a poroviscoelastic fibril-reinforced finite element study,” Journal of biomechanics, vol. 37, no. 3, pp. 357–366, 2004.

[66] W. Wilson, C. Van Donkelaar, B. Van Rietbergen, and R. Huiskes, “A fibril-reinforced poroviscoelastic swelling model for articular cartilage,” Journal of biomechanics, vol. 38, no. 6, pp. 1195–1204, 2005.

[67] E. A. Makris, P. Hadidi, and K. A. Athanasiou, “The knee meniscus: structure–function, pathophysiology, current repair techniques, and prospects for regeneration,” Biomaterials, vol. 32, no. 30, pp. 7411–7431, 2011.

[68] P. Atkinson, T. Atkinson, C. Huang, and R. Doane, “A comparison of the mechanical and dimensional properties of the human medial and lateral patellofemoral ligaments,” in Proceedings of the 46th Annual Meeting of the Orthopaedic Research Society, Orlando, FL, 2000.

[69] L. Blankevoort and R. Huiskes, “Ligament-bone interaction in a three-dimensional model of the knee,” Journal of biomechanical engineering, vol. 113, no. 3, pp. 263–269, 1991.

[70] L. Schatzmann, P. Brunner, and H. Stäubli, “Effect of cyclic preconditioning on the tensile properties of human quadriceps tendons and patellar ligaments,” Knee Surgery, Sports Traumatology, Arthroscopy, vol. 6, no. 1, pp. S56–S61, 1998.

[71] D. F. Villegas, J. A. Maes, S. D. Magee, and T. L. H. Donahue, “Failure properties and strain distribution analysis of meniscal attachments,” Journal of biomechanics, vol. 40, no. 12, pp. 2655–2662, 2007.

[72] T. C. Pataky, “Generalized n-dimensional biomechanical field analysis using statistical parametric mapping,” Journal of biomechanics, vol. 43, no. 10, pp. 1976–1982, 2010.

[73] M. Adouni and A. Shirazi-Adl, “Partitioning of knee joint internal forces in gait is dictated by the knee adduction angle and not by the knee adduction moment,” Journal of biomechanics, vol. 47, no. 7, pp. 1696–1703, 2014.

[74] H. Marouane, A. Shirazi-Adl, and M. Adouni, “Alterations in knee contact forces and centers in stance phase of gait: A detailed lower extremity musculoskeletal model,” Journal of biomechanics, vol. 49, no. 2, pp. 185–192, 2016.

[75] M. Adouni and A. Shirazi-Adl, “Evaluation of knee joint muscle forces and tissue stresses-strains during gait in severe oa versus normal subjects,” Journal of orthopaedic research, vol. 32, no. 1, pp. 69–78, 2014.

[76] D. Kumar, K. T. Manal, and K. S. Rudolph, “Knee joint loading during gait in healthy controls and individuals with knee osteoarthritis,” Osteoarthritis and cartilage, vol. 21, no. 2, pp. 298–305, 2013.

[77] S. C. O’reilly, A. Jones, K. R. Muir, and M. Doherty, “Quadriceps weakness in knee osteoarthritis: the effect on pain and disability,” Annals of the rheumatic diseases, vol. 57, no. 10, pp. 588–594, 1998.

[78] P. O. Bolcos, M. E. Mononen, A. Mohammadi, M. Ebrahimi, M. S. Tanaka, M. A. Samaan, R. B. Souza, X. Li, J.-S. Suomalainen, J. S. Jurvelin et al., “Comparison between kinetic and kinetic-kinematic driven knee joint finite element models,” Scientific reports, vol. 8, no. 1, pp. 1–11, 2018.

[79] D. J. Saxby, B. A. Killen, C. Pizzolato, C. Carty, L. Diamond, L. Modenese, J. Fernandez, G. Davico, M. Barzan, G. Lenton et al., “Machine learning methods to support personalized neuromusculoskeletal modelling,” Biomechanics and Modeling in Mechanobiology, vol. 19, no. 4, pp. 1169–1185, 2020.

[80] C. Pizzolato, V. B. Shim, D. G. Lloyd, D. Devaprakash, S. J. Obst, R. Newsham-West, D. F. Graham, T. F. Besier, M. H. Zheng, and R. S. Barrett, “Targeted achilles tendon training and rehabilitation using personalized and real-time multiscale models of the neuromusculoskeletal system,” Frontiers in Bioengineering and Biotechnology, vol. 8, p. 878, 2020.

[81] B. Killen, S. B. da Luz, D. Lloyd, A. Carleton, J. Zhang, T. Besier, and D. Saxby, “Automated creation and tuning of personalised muscle paths for opensim musculoskeletal models of the knee joint,” Biomechanics and Modeling in Mechanobiology, pp. 1–13, 2020.

[82] Z. I. Nejad, K. Khalili, S. H. H. Nasab, P. Schütz, P. Damm, A. Trepczynski, W. R. Taylor, and C. R. Smith, “The capacity of generic musculoskeletal simulations to predict knee joint loading using the cams-knee datasets,” Annals of biomedical engineering, vol. 48, no. 4, pp. 1430–1440, 2020.

[83] J. P. Walter, A. L. Kinney, S. A. Banks, D. D. D’Lima, T. F. Besier, D. G. Lloyd, and B. J. Fregly, “Muscle synergies may improve optimization prediction of knee contact forces during walking,” Journal of biomechanical engineering, vol. 136, no. 2, 2014.

[84] J. L. Silverberg, S. Dillavou, L. Bonassar, and I. Cohen, “Anatomic variation of depth-dependent mechanical properties in neonatal bovine articular cartilage,” Journal of Orthopaedic Research, vol. 31, no. 5, pp. 686–691, 2013.

[85] S. Treppo, H. Koepp, E. C. Quan, A. A. Cole, K. E. Kuettner, and A. J. Grodzinsky, “Comparison of biomechanical and biochemical properties of cartilage from human knee and ankle pairs,” Journal of Orthopaedic Research, vol. 18, no. 5, pp. 739–748, 2000.

[86] H. N. Beidokhti, D. Janssen, S. van de Groes, J. Hazrati, T. Van den Boogaard, and N. Verdonschot, “The influence of ligament modelling strategies on the predictive capability of finite element models of the human knee joint,” Journal of biomechanics, vol. 65, pp. 1–11, 2017.

[87] L. P. Räsänen, P. Tanska, Š. Zbỳň, C. C. van, S. Trattnig Donkelaar, M. T. Nieminen, and R. K. Korhonen, “The effect of fixed charge density and cartilage swelling on mechanics of knee joint cartilage during simulated gait,” Journal of biomechanics, vol. 61, pp. 34–44, 2017.

[88] L. P. Räsänen, M. E. Mononen, M. T. Nieminen, E. Lammentausta, J. S. Jurvelin, R. K. Korhonen, and O. Investigators, “Implementation of subject-specific collagen architecture of cartilage into a 2d computational model of a knee joint—data from the osteoarthritis initiative (oai),” Journal of orthopaedic research, vol. 31, no. 1, pp. 10–22, 2013.

[89] L. P. Räsänen, P. Tanska, M. E. Mononen, E. Lammentausta, Š. Zbỳň, M. S. Venäläinen, P. Szomolanyi, C. C. van Donkelaar, J. S. Jurvelin, S. Trattnig et al., “Spatial variation of fixed charge density in knee joint cartilage from sodium mri–implication on knee joint mechanics under static loading,” Journal of biomechanics, vol. 49, no. 14, pp. 3387–3396, 2016.

[90] L. P. Räsänen, M. E. Mononen, E. Lammentausta, M. T. Nieminen, J. S. Jurvelin, and R. K. Korhonen, “Three dimensional patient-specific collagen architecture modulates cartilage responses in the knee joint during gait,” Computer methods in biomechanics and biomedical engineering, vol. 19, no. 11, pp. 1225–1240, 2016.

[91] J. Y. Bae, K. S. Park, J. K. Seon, I. Jeon, E. K. Song et al., “Biomechanical analysis of the effects of medial meniscectomy on degenerative osteoarthritis,” Medical & biological engineering & computing, vol. 50, no. 1, pp. 53–60, 2012.

