## Supplementary materials for "An EMG-assisted Muscle-Force Driven Finite Element Analysis Pipeline to Investigate Joint- and Tissue-Level Mechanical Responses in Functional Activities: Towards a Rapid Assessment Toolbox"

### 1. Method

#### 1.1. Subjects

Fifteen subjects (6 male and 9 female) were recruited in this study. Table S.1 shows the participants' characteristics. The inclusion criteria were to have KOA according to the KOA clinical definition (i.e., the existence of both pain and an evident radiographic joint tissue deterioration) [1] and knee pain on most of the days during the last month. The exclusion criteria were the existence of any record of lower limb surgeries or disorders such as ligament or tendon rupture or presence of pain in any other body parts except for the knee. The more affected leg of each subject (in terms of the clinically diagnosed KOA and also the pain score) was selected and was anchored for that subject during this study (i.e. a total of 15 knee joints, one knee per each subject). All the procedures were approved by the ethics committee of the Hospital District of Northern Savo (permission No. 750/2018), and written informed consent was obtained from each subject.

**Table S1.** Participants' characteristics.

| | **Mean ± SD** | **Range** |
|---|---|---|
| **Age (years)** | 62.4 ± 7.8 | 51.5 - 73.2 |
| **Height (m)** | 1.70 ± 0.10 | 1.55 - 1.85 |
| **Body weight (kg)** | 84.4 ± 19.1 | 59.0 - 122.3 |
| **Body mass index (kg/m2)** | 29.3 ± 6.8 | 20.5 - 46.8 |
| **Pain at the affected knee during the last month (VAS)** | 40.7 ± 21.2 | 2.8 - 74 |
| **Habitual walking speed (m/s)** | 1.34 ± 0.14 | 1.01 – 1.62 |

#### 1.2. Daily activities and rehabilitation exercises

Seven daily activities consisted of walking (at a habitual speed of 1.34 ± 0.14 m/s and at a constant speed of 20 ± 0.05 m/s), picking up a light-weight object from the ground, stair ascending and descending, and chair sit-to-stand and stand-to-sit. First, we asked the subjects to perform six trials of 15-meters habitual walking. At each trial, the average walking speed over a 3-meter section (after a 7-meter run-up) was measured using photocells. Then, the fastest and slowest trials were excluded, and the habitual walking speed was determined as the average of the four remaining trials. Subjects walked with their own outfit

*

and shoes. Except for determining the habitual walking speed, subjects performed all the daily tasks and rehabilitation exercises with minimal clothing (spandex shorts and a shirt and barefoot).

Subjects performed daily activities in random order, and five successful repetitions of each activity were recorded. The collected motion data consisted of marker trajectories (100 Hz, Vicon, UK), ground reaction forces (two force plates, 1000 Hz, OR6-7MA, AMTI, USA), and EMGs (1000 Hz, Biomonitor ME6000, Bittium Biosignals, Finland). Ground reaction forces in the stair ascent and descent were recorded using the floor-mounted force plates and a custom-made staircase. The staircase consisted of two steps (15 cm and 30 cm high) and a platform (45 cm high), The 30 cm high step was placed on a force plate and was mechanically isolated from the surroundings. EMGs from 8 muscles of the leg of interest including vastus medialis and lateralis, rectus femoris, medial and lateral gastrocnemius, biceps femoris, semitendinosus, and gluteus medius were recorded according to the SENIAM instructions [2]. In addition to the mentioned measurements, magnetic resonance images (MRI) were taken from the knee of interest of each subject, using a 0.18 T scanner (Esaote E-Scan XQ, Esaote, Genoa, Italy). MRIs consisted of the three dimensional (3D CE, with 0.89 mm slice thickness and 0.625 in-plane resolution), Spin Echo T2 (with 3.5 mm slice thickness and 0.78 mm in-plane resolution), and STIR (4 mm slice thickness and 0.78 mm in-plane resolution) sequences.

Marker trajectories and GRFs were filtered using a second-order zero-lag Butterworth low-pass filter with cut-off frequencies of 6 Hz and 30 Hz, respectively. Utilizing the MOtoNMS [3], EMG signals were band-pass filtered (30-300 Hz), rectified, low-pass filtered (6 Hz), and in the end, normalized against all the dynamic trials of the subject as well as maximum isometric voluntary contractions [4]. The maximum isometric voluntary contraction trials were conducted for hip abduction/adduction, hip flexion/extension, knee flexion/extension, and ankle dorsi/plantar flexion.

#### 1.3. FE model

##### 1.3.1. Material models

A fibril-reinforced poroviscoelastic (FRPVE) material model [5,6] was utilized for the cartilages and the depth-dependent Benninghoff-type (arcade) architecture of collagen fibers was implemented as split-lines for femoral, tibial, and patellar cartilages [7–10]. Menisci were modeled as a fibril-reinforced poroelastic (FRPE) material [5,11]. More details about the implementation of the FRPVE material model can be found from our previous studies [12]. The total stress in the FRP(V)E model ($\sigma_t$) consists of the non-fibrillar matrix stress ($\sigma_{nf}$), collagen fibril stress ($\sigma_f$), and fluid pressure ($p$):

$$\sigma_t = \sigma_{nf} + \sigma_f - p\mathbf{I} \tag{1}$$

where $\mathbf{I}$ is the unit tensor. The non-fibrillar matrix was modeled by compressible neo-Hookean properties. The stress within the non-fibrillar matrix is given by [6]:

$$\sigma_{nf} = \frac{1}{2}K(J - \frac{1}{J})\mathbf{I} + \frac{G}{J}(\mathbf{F}.\mathbf{F}^T - J^{(2/3)}\mathbf{I}) \tag{2}$$

$$K = \frac{E_{nf}}{3(1 - 2\nu_{nf})} \tag{3}$$

$$G = \frac{E_{nf}}{2(1 + \nu_{nf})} \tag{4}$$

where $G$ and $K$ are the shear and bulk moduli of the non-fibrillar matrix, $J$ is the determinant of the deformation tensor $\mathbf{F}$, $E_{nf}$ and $\nu_{nf}$ are Young's modulus and Poisson's ratio of the non-fibrillar matrix, respectively. A strain-dependent permeability ($k$) [13] is given by:

$$k = k_0(\frac{1+e}{1+e_0})^M \tag{5}$$

where $k_0$ is the initial permeability, $e$ and $e_0$ are the current and the initial void ratios, and $M$ is a positive constant. The fluid fraction was assumed to be depth-dependent in equilibrium [14] as:

$$n_{(f,eq)} = 0.85 - 0.15d_n \tag{6}$$

where $d_n$ is the normalized depth (0 at the surface and 1 at the cartilage-bone interface).

The cartilage collagen fibers were modeled as a viscoelastic material. In the material model, a nonlinear spring (with the strain-dependent modulus $E_\epsilon \epsilon_f$) is in series with a linear dashpot (with the damping coefficient $\eta$). This nonlinear spring-dashpot system is in parallel with a linear spring (with the initial modulus $E_0$). Fibrils were assumed to only resist tension; thus, the collagen fibril stress of the cartilage was formulated as [6,15]:

$$\sigma_f = \begin{cases} -\frac{\eta}{\sqrt{(\sigma_f - E_0\epsilon_f)E_\epsilon}}\dot{\sigma}_f + E_0\epsilon_f + (\eta + \frac{\eta E_0}{\sqrt{(\sigma_f - E_0\epsilon_f)E_\epsilon}})\dot{\epsilon}_f & , \quad \epsilon_f > 0 \\ 0 & , \quad \epsilon_f \leq 0 \end{cases} \tag{7}$$

where $\sigma_f$ and $\epsilon_f$ are the fibril stress and strain, and $\dot{\sigma}_f$ and $\dot{\epsilon}_f$ are the fibril stress and strain rates.

The collagen fibers within the menisci were modeled as linear elastic (with Young's modulus of $E_f$). The menisci collagen fiber stress was formulated as [16]:

$$\sigma_f = \begin{cases} E_f\epsilon_f & , \quad \epsilon_f > 0 \\ 0 & , \quad \epsilon_f \leq 0 \end{cases} \tag{8}$$

The collagen fiber network consisted of primary and secondary fibrils [6]. The primary collagen fibrils form a depth-dependent arcade-like structure [17], while the secondary fibrils are randomly organized in 13 different random orientations [6]. Secondary fibrils mainly replicate the inter-fibril connections and cross-links in the collagen network. Consequently, defining $C$ as the amount of the primary fibrils with respect to the secondary fibrils, the stresses are given by [6]:

$$\begin{cases} \sigma_{f,p} = C\sigma_f \\ \sigma_{f,s} = \sigma_f \end{cases} \tag{9}$$

**Table S2.** Material parameters for the knee joint cartilages and menisci

| **Material parameter** | **Femoral cartilage** | **Tibial cartilage** | **Patellar cartilage** | **Menisci** |
|---|---|---|---|---|
| $E_{nf}(MPa)$ | 0.215 | 0.106 | 0.505 | 0.5 |
| $v_{nf}(-)$ | 0.15 | 0.15 | 0.15 | 0.36 |
| $k_0(\frac{m^4}{N.s} \times 10^{-15})$ | 6 | 18 | 1.9 | 1.25 |
| $\eta(MPa.s)$ | 1062 | 1062 | 1062 | - |
| $E_0(MPa)$ | 0.92 | 0.18 | 1.88 | - |
| $E_f(MPa)$ | - | - | - | 28 |
| $E_\epsilon(MPa)$ | 150 | 23.6 | 597 | - |
| C(-) | 12.16 | 12.16 | 12.16 | 12.16 |
| $M(-)$ | 5.09 | 15.64 | 15.93 | 5.09 |
| $n_{f,eq}(-)$ | $0.85 - 0.15d_n$ | $0.85 - 0.15d_n$ | $0.85 - 0.15d_n$ | 0.72 |

Consistent with our previous study [18], ligaments and tendons were modelled as spring bundles to have sufficient accuracy in the estimated parameters while keeping the computational demand reasonable. Ligament and tendon insertion points were segmented from the template MRIs. Non-linear spring bundles were used to replicate the Anterior cruciate ligament (ACL, 60 springs), posterior cruciate ligament (PCL, 60 springs), lateral collateral ligament (LCL, 18 springs), and medial collateral ligament (MCL, 18 springs). Utilizing a bundle of springs provides the ligament model with compression-tension nonlinearity with different properties along and perpendicular to the fibril/spring directions. The slack, toe, and linear regions of the ligaments were formulated according to the study by Blankevoort et al. [19] as:

$$f_s = \begin{cases} 0 & , \quad \epsilon_s < 0 \\ \frac{1}{4}K_s\epsilon_s^2/\epsilon_l & , \quad 0 \leq \epsilon_s \leq 2\epsilon_l \\ K_s(\epsilon_s - \epsilon_l) & , \quad \epsilon_s \geq 2\epsilon_l \end{cases} \tag{10}$$

where $f_s$ is the tensile force in each ligament element, $K_s$ is the ligament stiffness [19], $\epsilon_l$ represents the end of the toe region and was set to 0.03 [20], and $\epsilon$ is the current strain in the ligament.

The medial and lateral patellofemoral ligaments (MPFL and LPFL, respectively) were modelled using linear spring bundles with no compressive resistance. The spring stiffness ( i.e. as the bundle) was defined as 15.9 N/mm for MPFL and 11.7 N/mm for LPFL [21]. Menisci horn attachments were modelled as linear spring bundles with a total stiffness of 336 N/mm and 381 N/mm for anterior and posterior sides, respectively [22]. Similarly, the patellar tendon was represented by two springs (no resistance in compression) with a total spring constant equal to 545 N/mm [23].

##### 1.3.2. The atlas-based FE modeling toolbox (morphometry and FE model scaling)

The morphometry from the MRIs for the FE model scaling consisted of the following measurements. In the sagittal plane, the height and thickness of the patella (from the middle of the trochlear groove), the maximum anteroposterior distance for the ellipse-like shaped medial and lateral femoral condyles, and the maximum outer distance of medial and lateral menisci were measured [24] (Fig. 2). In the coronal plane, first, the slice with the maximum width of the femoral condyles was selected (Fig. 2). Then from the selected slice, the width of the femoral and tibial cartilages, the outer distance between medial and lateral menisci, the thickness of the medial and lateral menisci, and medial and lateral tibiofemoral joint space were measured (Fig. 2). The width of the patella was measured from the slice in the transverse plane with the maximum femoral width. A musculoskeletal radiologist supervised all the morphometry to assure consistency between the measurements for different subjects.

Following the morphometry, different parts of the template was scaled as follows. The anteroposterior dimension of the femur was linearly scaled using the average of medial and lateral anteroposterior measures. The mediolateral dimension of the femur was scaled using the corresponding measure. likewise, the medial and lateral menisci and tibia were each scaled based on the corresponding medial and lateral dimensions. Thickness of the femoral and tibial cartilage were scaled according to the measured tibiofemoral joint space (Fig. 2). Patella was scaled using the its measured thickness, height, and width (Fig. 2). Ligament insertion points in the femur, tibia, and patellar were scaled according to the corresponding morphometry of the femur, tibia (average of medial and lateral plateaus), and patella. Within MATLAB, and using the STL format of the scaled geometries, the femur and menisci were moved upward, and the patella was moved anteriorly (with 1mm increments) to automatically find the best assembled alignment within the FE model parts (i.e., femur, tibia, patella, and menisci) with a minimum gap between the parts and no contact overclosure.

##### 1.3.3. FE model loading and boundary conditions

Loading and boundary condition inputs to the FE models obtained from the MS models of the study consisted of 1) knee flexion angle, 2) knee abduction/adduction and internal/external moments, 3) abduction/adduction and internal/external moments generated by the muscles around the tibiofemoral joint, 4) flexion/extension, abduction/adduction, and internal/external moments generated by the quadriceps muscles around the patellofemoral joint, 5) total tibiofemoral JCF, and 6) total patellofemoral JCFs (Fig. 1).

The FE model coordinate system was defined similar to the tibial coordinate system of the OpenSim, and was fixed to the tibia. The knee flexion was estimated from OpenSim inverse kinematics (IK) and the knee abduction/adduction and internal/external moment were calculated utilizing OpenSim inverse dynamics (ID). OpenSim calculates both IK and ID results in the tibial coordinate system. Muscle moment arms were extracted within the muscle analysis toolbox of the OpenSim. We should notice that the abduction/adduction and internal/external degrees of freedom were added to the tibiofemoral joint mechanism (but they were locked) to calculate knee abduction/adduction and internal/external moments and corresponding muscle moment arms. Similarly, flexion/extension, abduction/adduction, and internal/external DoFs were added (and locked) to the patellofemoral joint to be able to calculate muscle moment arms around the patella. The muscle moment arms were extracted for patella flexion/extension, abduction/adduction, and internal/external DoFs.

In our previous muscle-force driven FE model implementation [18,25], the muscle forces were first subtracted from the total JCF, and then were applied into the FE models on their insertion points along with their effective line of action. We developed this approach to include the moments generated by the muscles around the knee joint into the FE model, consistent with the MS model. However, this implementation requires an OpenSim plugin [26], which brings difficulties such as compatibility issues with different OpenSim versions. Also, excessive efforts are needed to implement muscle insertion points and muscle lines of action in each FE model. Therefore, in this study, we redesigned the muscle-force implementation in the FE models to facilitate the interconnection of the MS and FE models within an automated pipeline.

The moment generated by each muscle with respect to the reference point in either femur or patella was calculated by multiplying the muscle moment arms by the muscle force. The muscle forces were used from either static-optimization outputs or CEINMS results, correspondingly. The muscle moment around the tibiofemoral joint was calculated in the tibial coordinate system. However, the muscle moments around the patellofemoral joint were calculated in the patella coordinate system, since OpenSim reports those moment arms in the patellar coordinate system. Thus, the calculated muscle moments around the patellofemoral joint were transformed into the tibial coordinate system within an in-house MATLAB script and according to the IK results.

The JCFs were calculated using the joint analysis toolbox of the OpenSim. The tibiofemoral JCF was calculated in the tibial coordinate system directly in OpenSim. However, the patellofemoral JCF was extracted in the patella coordinate system of the OpenSim, and then were transformed into the tibial coordinate system with the in-house script. This was done since OpenSim cannot report the patellofemoral JCF in the tibial coordinate system with the model used in the current study. More explanations about the coordinate systems, transformations, etc., can be found in our previous study [18].

All the nodes on the bottom of the tibia were fixed in the FE models. Consequently, inputs were applied to the femur and patella reference points. The only kinematic input to the FE model was knee flexion angle, and the rest of the DoFs (3 translational + 2 rotational DoFs of the femur, and 3 translational + 3 rotational DoFs of the patella) were set free. The femur reference point was defined at the center of rotation of the femur (i.e., the tibiofemoral joint center), and all the nodes located on the bone-cartilage interface of the femur were coupled to this reference point. Knee flexion angle, tibiofemoral JCF, and knee moments (including moments from ID and those generated by muscles) were applied to the femoral reference point. Similarly, a reference point was defined on the patella, and all the nodes located on the cartilage-subchondral bone interface were coupled to that reference point. Patellofemoral JCFs and moments generated by quadriceps muscles were applied to the patella reference point.

#### 1.4. The semi-autonomous pipeline

The whole workflow of the study (Fig. 1 and 2) was implemented in a semi-autonomous pipeline. Only the model scaling step (i.e., the MS models and the FE models) were performed manually. Although the MS scaling could be automated, it was done manually to be assured of precise scaling. Apart from that, the rest of the workflow was implemented by an in-house MATLAB script (version 2019b, MathWorks Inc., US) as follows:

1. The inverse kinematics, inverse dynamics, static optimization, JCF analysis, and muscle analysis were performed sequentially utilizing the OpenSim-MATLAB API.
2. Creating the CEINMS setup files (based on the scaled MS models), writing input files (i.e. uncalibrated model), running the calibration, and then executing the EMG-assisted MS analyses were performed with the MATLAB script.
3. Employing the OpenSim-MATLAB API, the results of the SO-based MS analysis and EMG-assisted MS analysis were read, and then the input files of the FE models were written by attaching the scaled FE models and the inputs to the FE models.
4. Then, the FE models were executed in Abaqus from MATLAB. After finishing the analysis, the FE results were extracted and processed using python scripts executed from MATLAB.
5. In the end, the results were organized in the SPSS format, and the statistical analysis was performed using SPSS syntax commands called from MATLAB.

It should be mentioned that all the results from each step (e.g., EMGs, IK, ID, JCF, etc.) were plotted and checked manually to ensure the outputs and the workflow.

### 2. Results

Complementary results of the study are illustrated in Figs. S1 to S18.

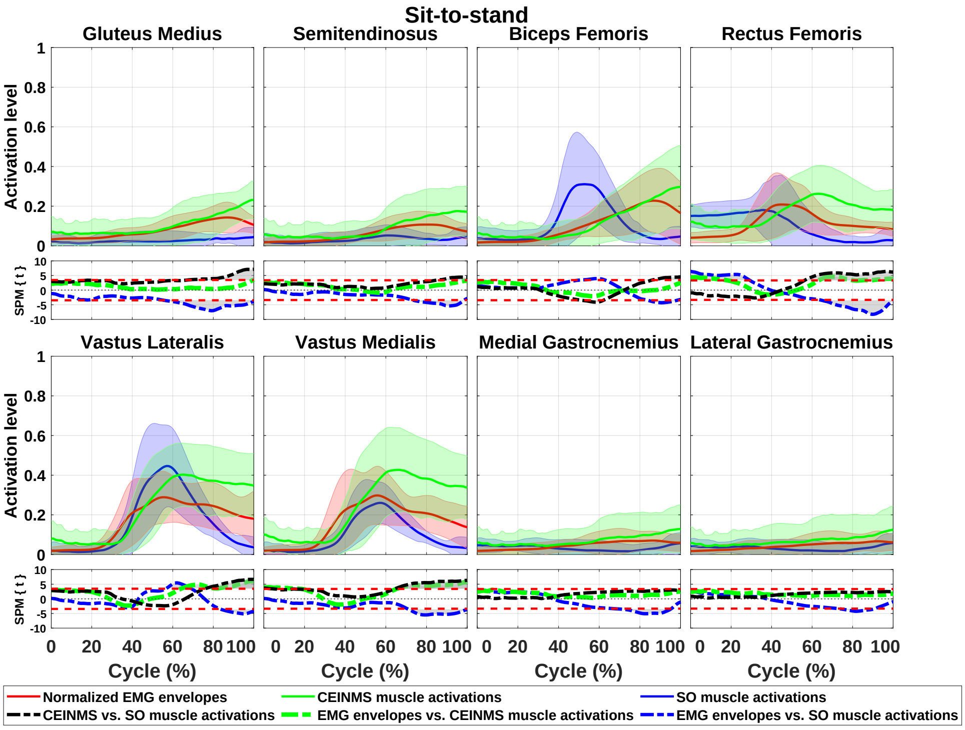


**Fig. S1:** Enveloped EMGs compared to muscle activations estimated by the SO-based and EMG-assisted MS models during sit-to-stand. The SPM plots (second and fourth rows) show comparisons between the enveloped EMGs and estimated muscle activations using paired sample t-test.

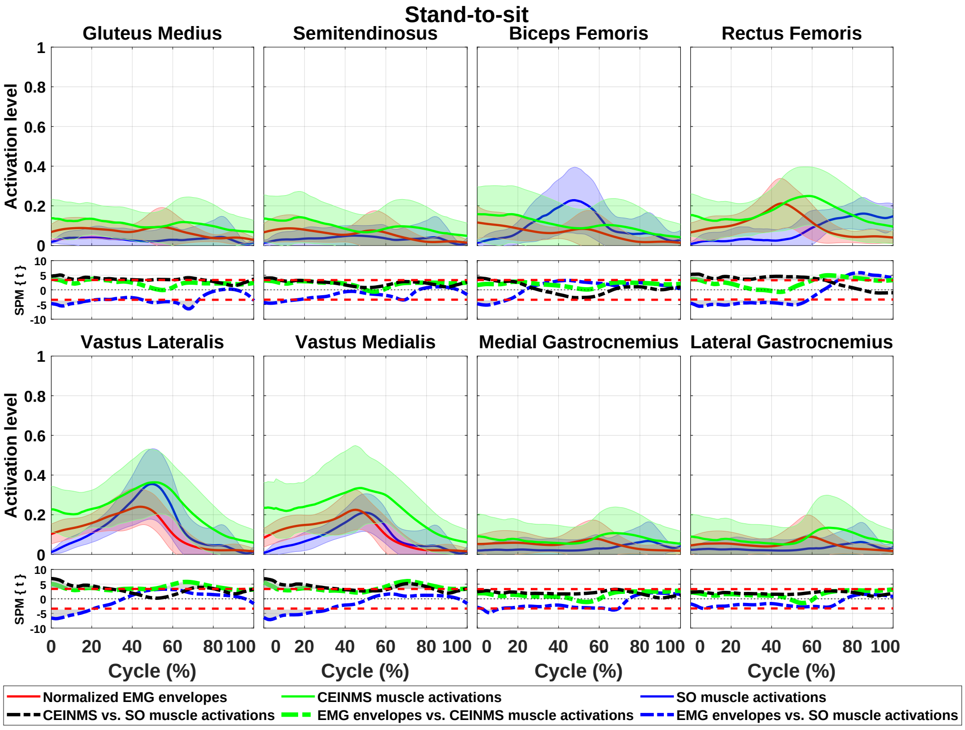


**Fig. S2:** Enveloped EMGs compared to muscle activations estimated by the SO-based and EMG-assisted MS models during stand-to-sit. The SPM plots (second and fourth rows) show comparisons between the enveloped EMGs and estimated muscle activations using paired sample t-test.

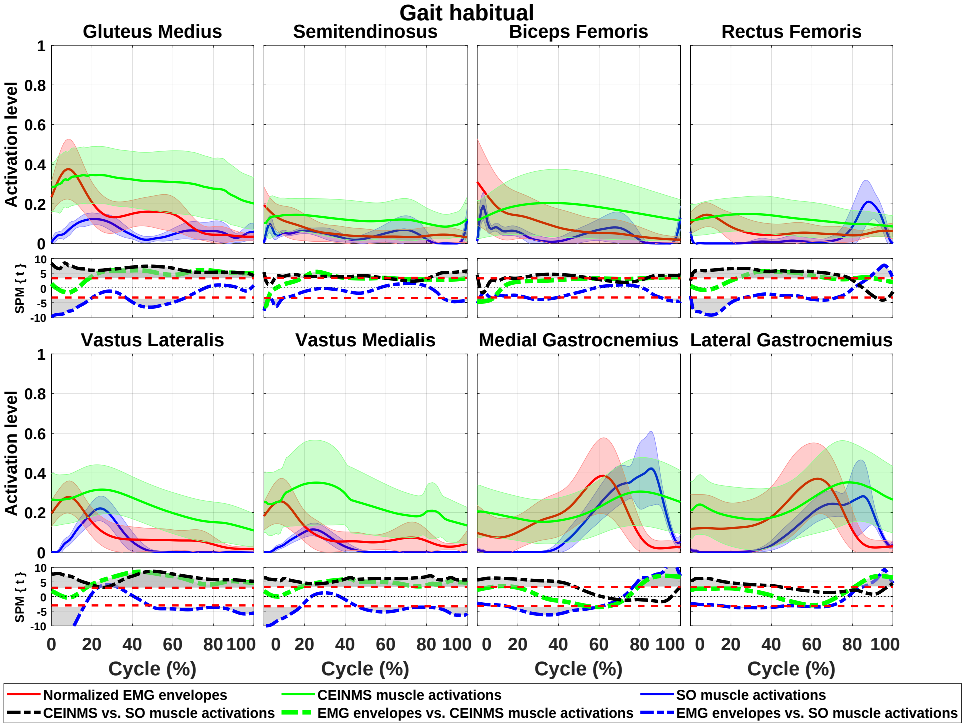


**Fig. S3:** Enveloped EMGs compared to muscle activations estimated by the SO-based and EMG-assisted MS models during gait habitual. The SPM plots (second and fourth rows) show comparisons between the enveloped EMGs and estimated muscle activations using paired sample t-test.

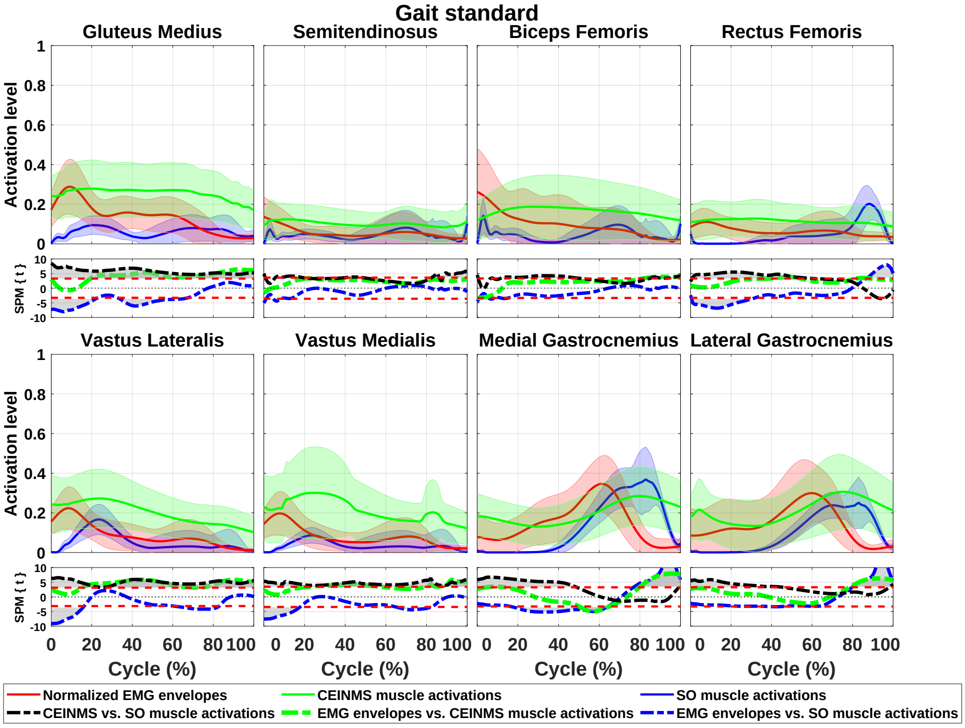


**Fig. S4:** Enveloped EMGs compared to muscle activations estimated by the SO-based and EMG-assisted MS models during gait standard. The SPM plots (second and fourth rows) show comparisons between the enveloped EMGs and estimated muscle activations using paired sample t-test.

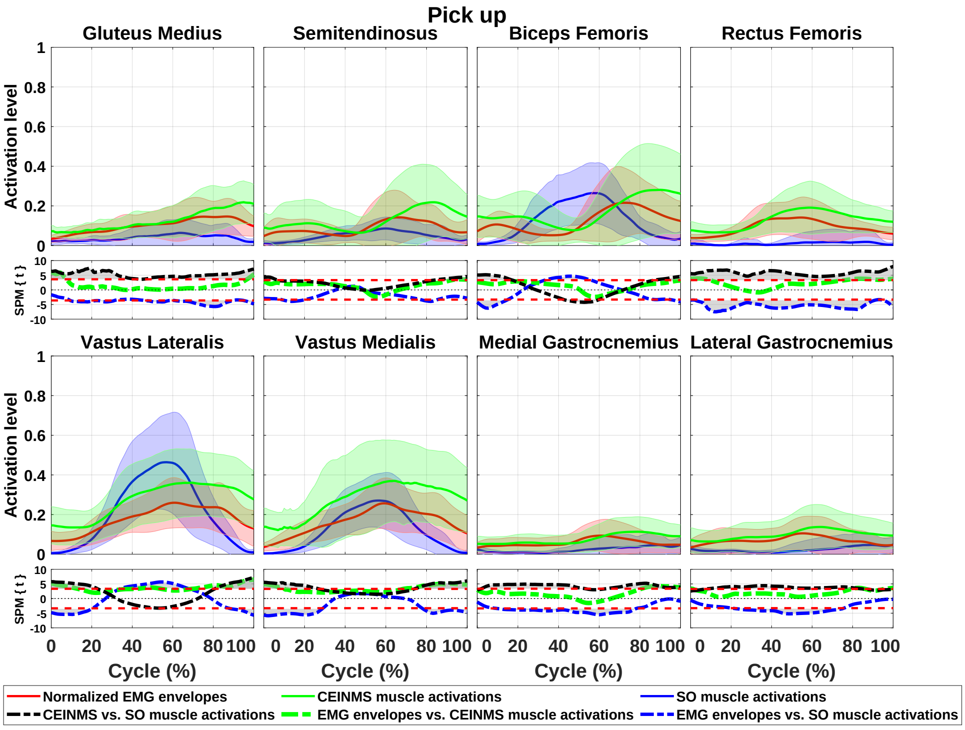


**Fig. S5:** Enveloped EMGs compared to muscle activations estimated by the SO-based and EMG-assisted MS models during pick up. The SPM plots (second and fourth rows) show comparisons between the enveloped EMGs and estimated muscle activations using paired sample t-test.

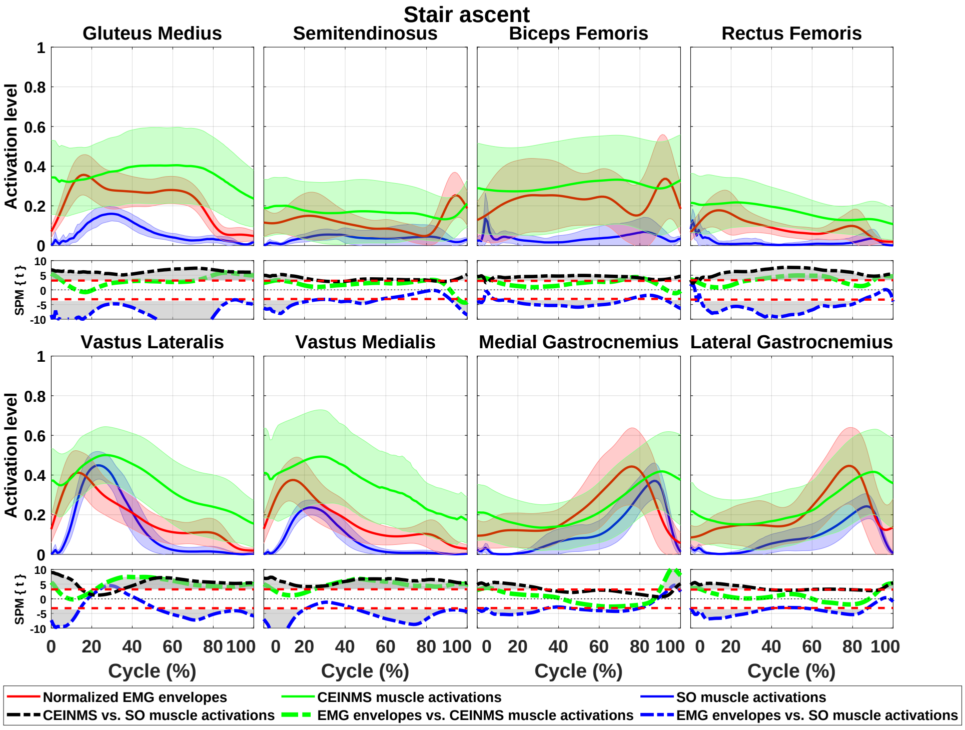


**Fig. S6:** Enveloped EMGs compared to muscle activations estimated by the SO-based and EMG-assisted MS models during stair ascent. The SPM plots (second and fourth rows) show comparisons between the enveloped EMGs and estimated muscle activations using paired sample t-test.

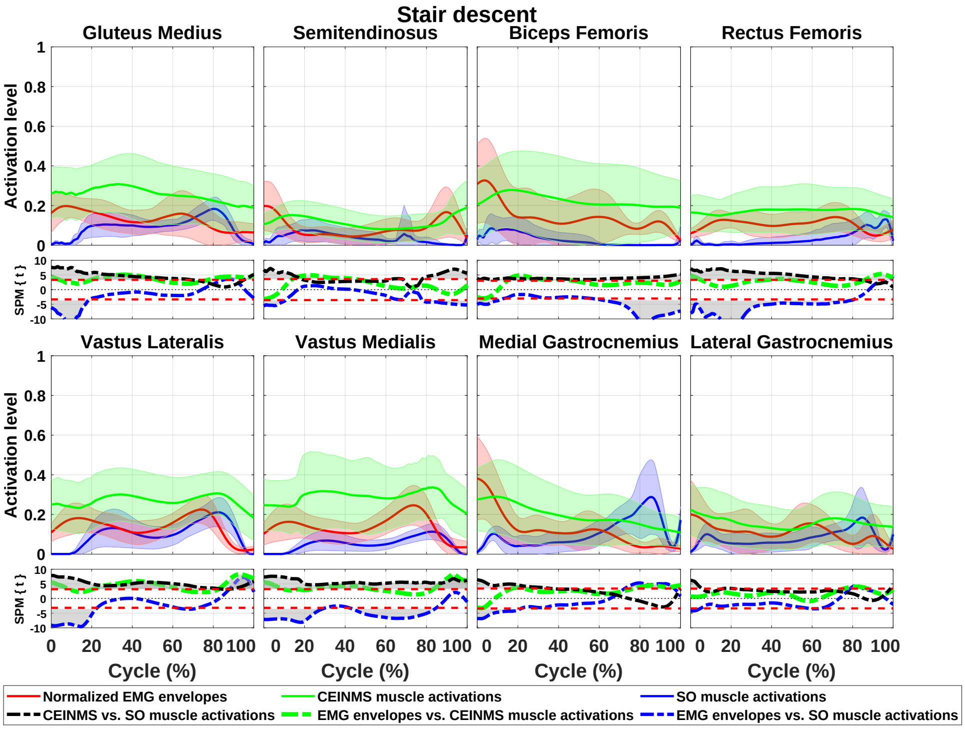


**Fig. S7:** Enveloped EMGs compared to muscle activations estimated by the SO-based and EMG-assisted MS models during stair descent. The SPM plots (second and fourth rows) show comparisons between the enveloped EMGs and estimated muscle activations using paired sample t-test.

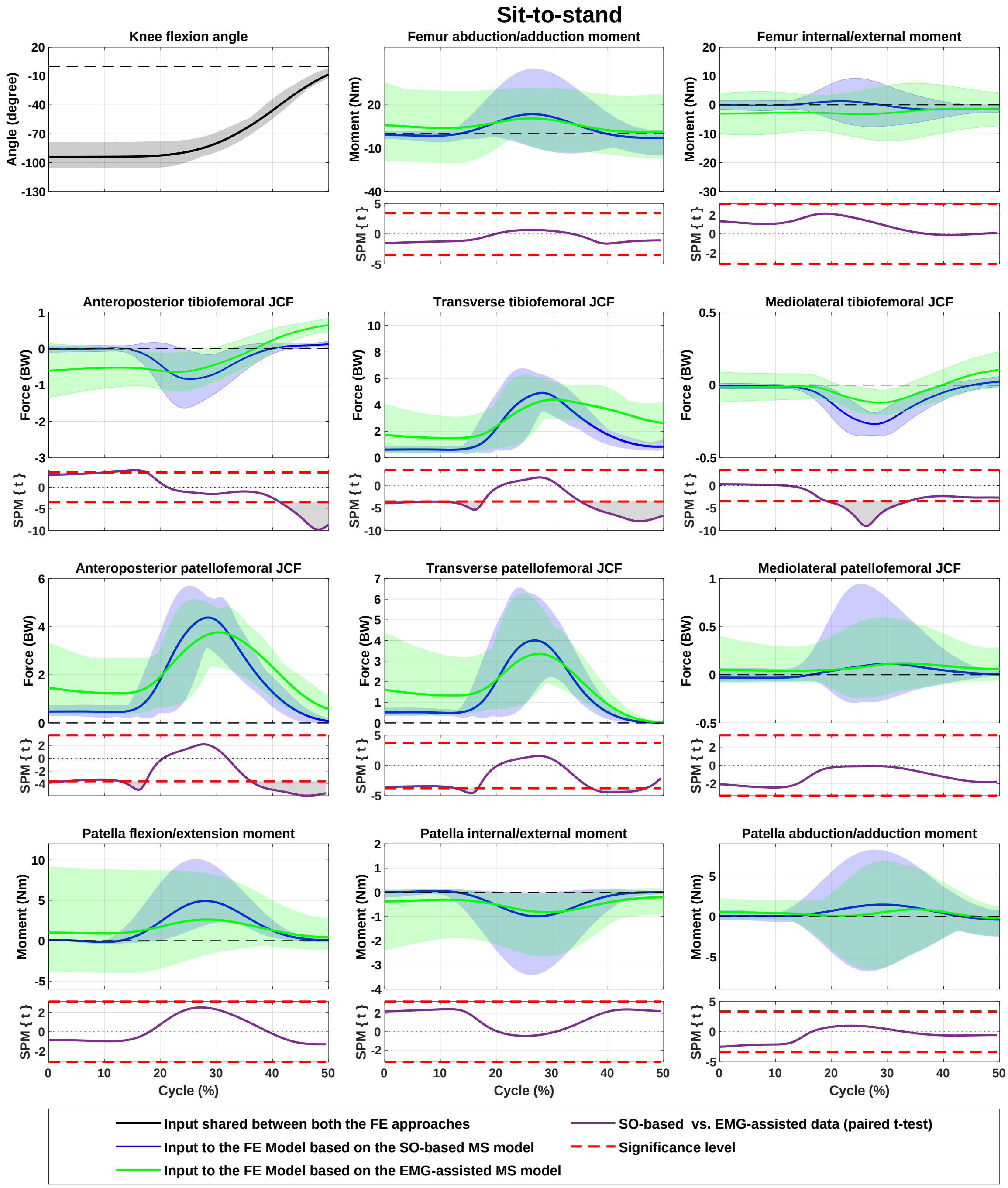


**Fig. S8:** Inputs to the FE models of the study during sit-to-stand, consisted of knee flexion angle, tibiofemoral joint moments (those calculated from inverse dynamics plus the moments generated by the muscles), tibiofemoral joint contact forces, patellofemoral joint contact forces, and patellofemoral moment generated by quadriceps muscles. Inputs estimated by the SO-based and EMG-assisted MS models are illustrated in blue and green, respectively. The SPM plots (in magenta) show comparisons between the results from the EMG-assisted and SO-based MS models, using paired sample t-test.

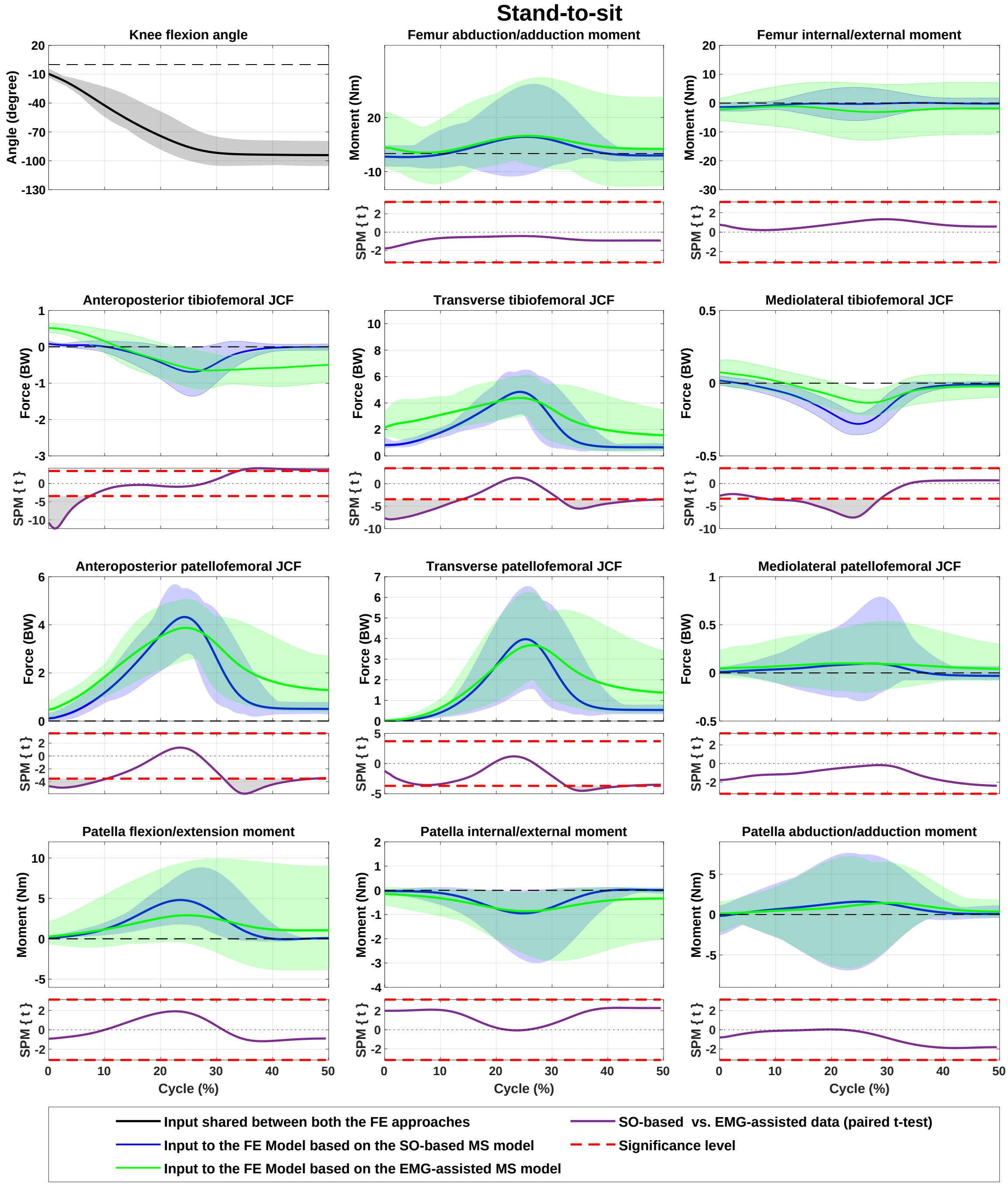


**Fig. S9:** Inputs to the FE models of the study during stand-to-sit, consisted of knee flexion angle, tibiofemoral joint moments (those calculated from inverse dynamics plus the moments generated by the muscles), tibiofemoral joint contact forces, patellofemoral joint contact forces, and patellofemoral moment generated by quadriceps muscles. Inputs estimated by the SO-based and EMG-assisted MS models are illustrated in blue and green, respectively. The SPM plots (in magenta) show comparisons between the results from the EMG-assisted and SO-based MS models, using paired sample t-test.

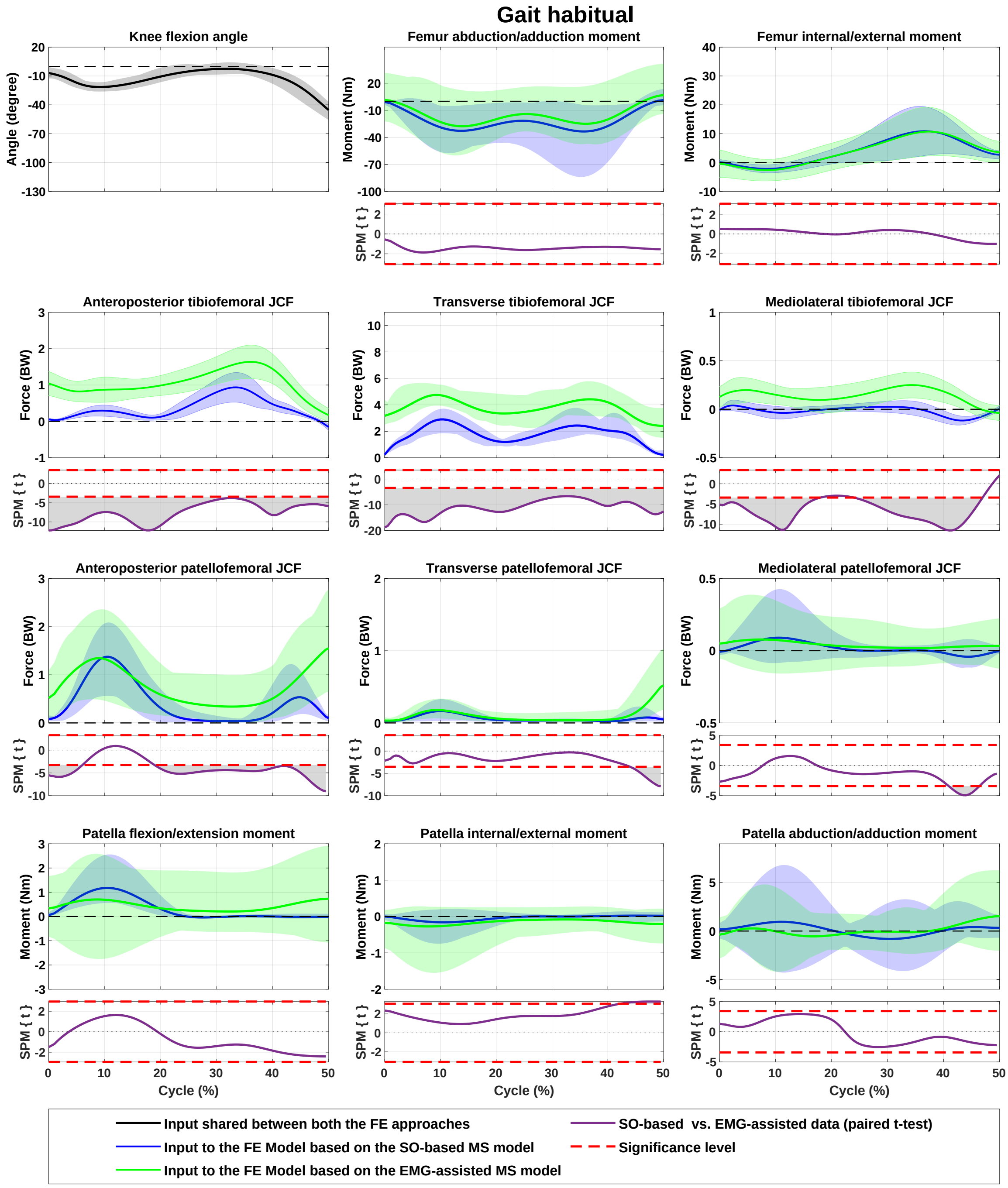


**Fig. S10:** Inputs to the FE models of the study during gait habitual, consisted of knee flexion angle, tibiofemoral joint moments (those calculated from inverse dynamics plus the moments generated by the muscles), tibiofemoral joint contact forces, patellofemoral joint contact forces, and patellofemoral moment generated by quadriceps muscles. Inputs estimated by the SO-based and EMG-assisted MS models are illustrated in blue and green, respectively. The SPM plots (in magenta) show comparisons between the results from the EMG-assisted and SO-based MS models, using paired sample t-test.

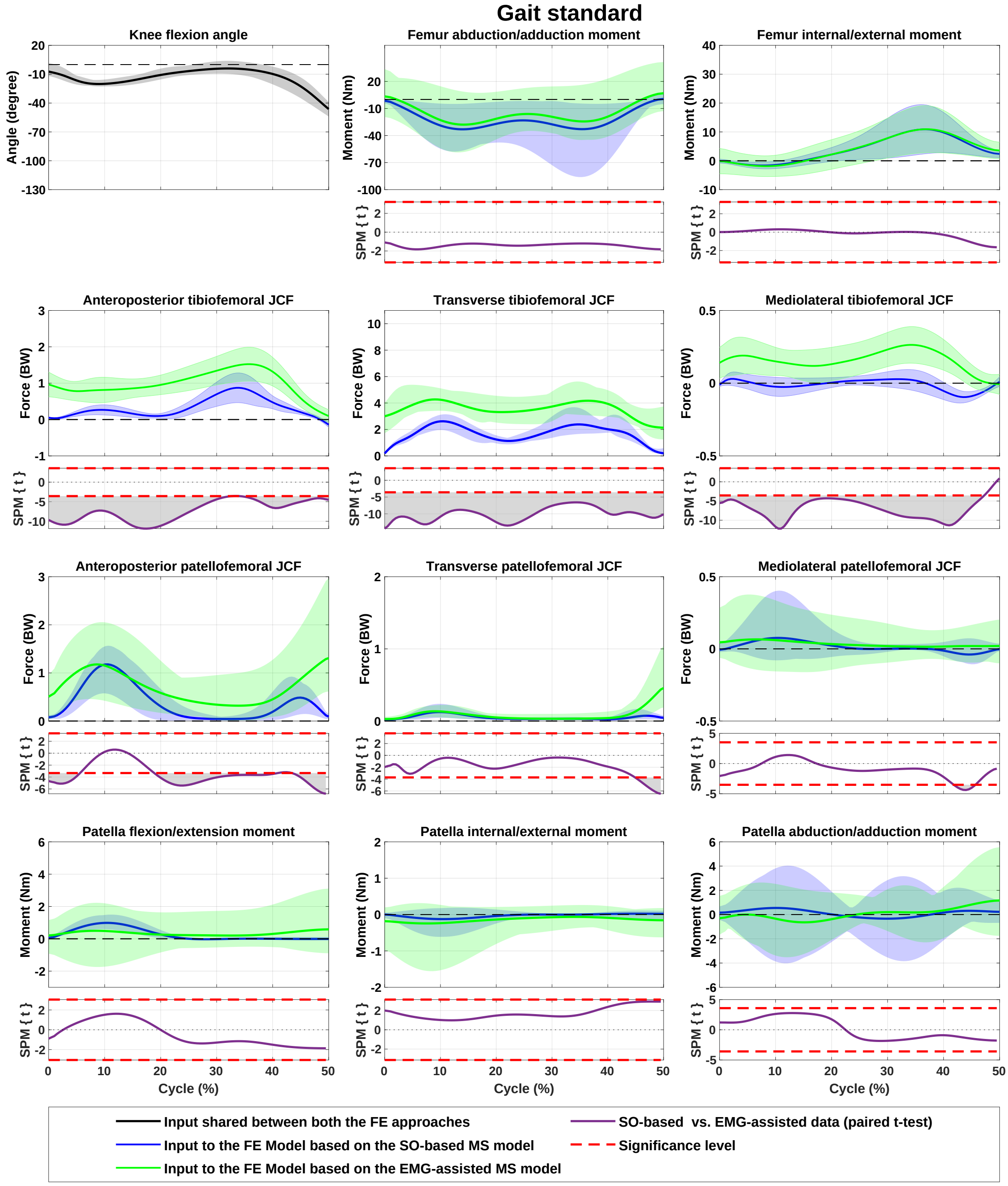


**Fig. S11:** Inputs to the FE models of the study during gait standard, consisted of knee flexion angle, tibiofemoral joint moments (those calculated from inverse dynamics plus the moments generated by the muscles), tibiofemoral joint contact forces, patellofemoral joint contact forces, and patellofemoral moment generated by quadriceps muscles. Inputs estimated by the SO-based and EMG-assisted MS models are illustrated in blue and green, respectively. The SPM plots (in magenta) show comparisons between the results from the EMG-assisted and SO-based MS models, using paired sample t-test.

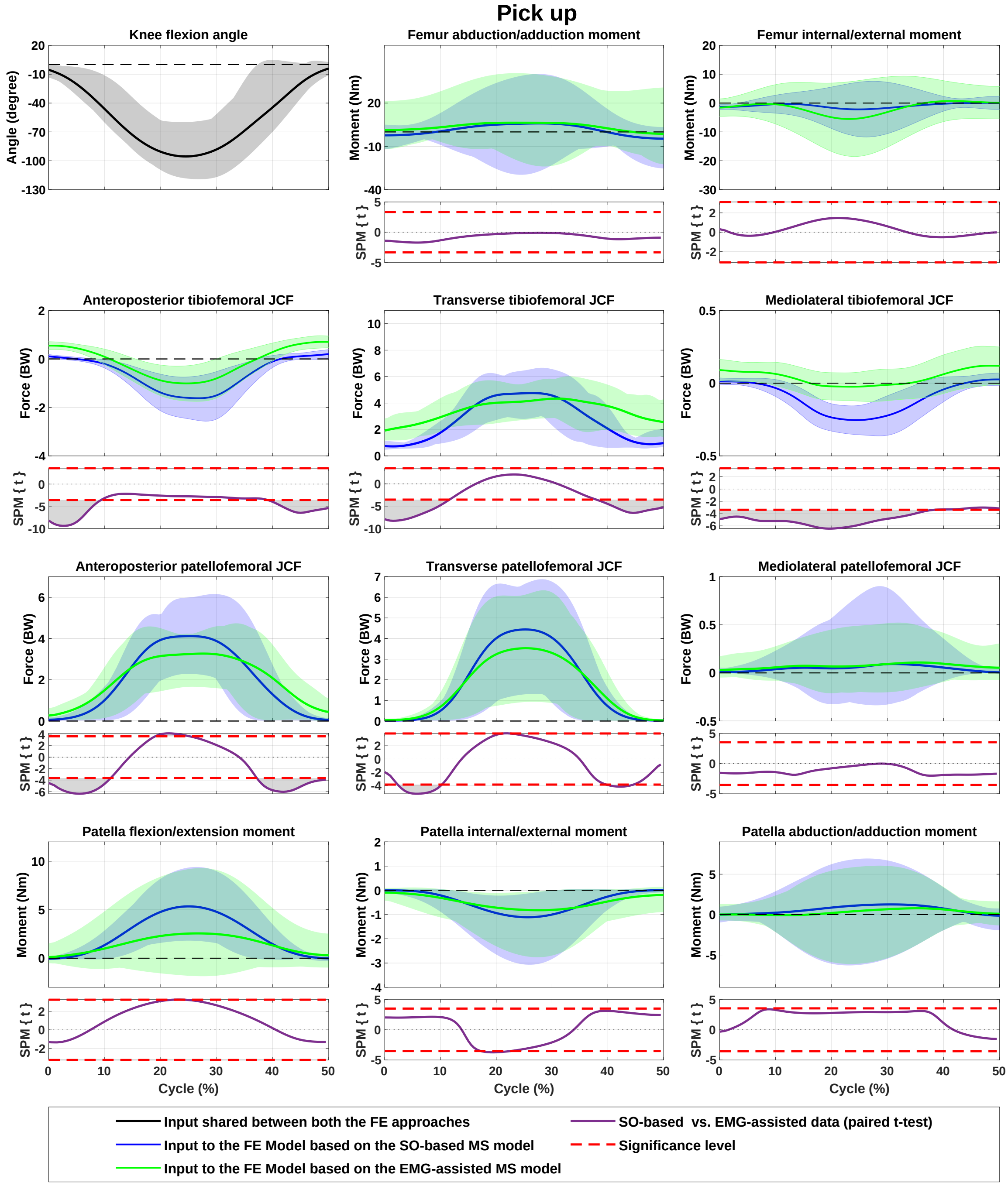


**Fig. S12:** Inputs to the FE models of the study during pick up, consisted of knee flexion angle, tibiofemoral joint moments (those calculated from inverse dynamics plus the moments generated by the muscles), tibiofemoral joint contact forces, patellofemoral joint contact forces, and patellofemoral moment generated by quadriceps muscles. Inputs estimated by the SO-based and EMG-assisted MS models are illustrated in blue and green, respectively. The SPM plots (in magenta) show comparisons between the results from the EMG-assisted and SO-based MS models, using paired sample t-test.

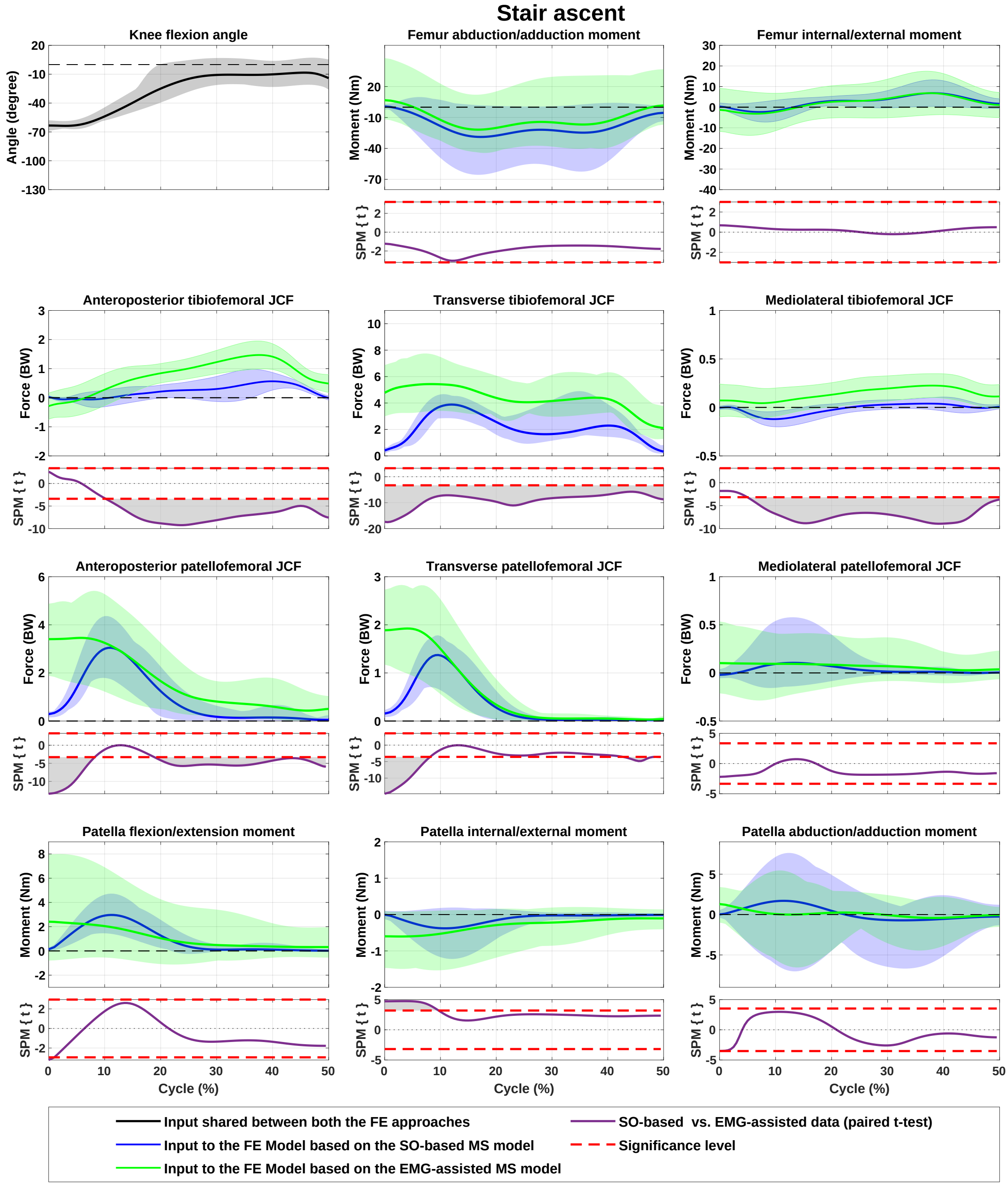


**Fig. S13:** Inputs to the FE models of the study during stair ascent, consisted of knee flexion angle, tibiofemoral joint moments (those calculated from inverse dynamics plus the moments generated by the muscles), tibiofemoral joint contact forces, patellofemoral joint contact forces, and patellofemoral moment generated by quadriceps muscles. Inputs estimated by the SO-based and EMG-assisted MS models are illustrated in blue and green, respectively. The SPM plots (in magenta) show comparisons between the results from the EMG-assisted and SO-based MS models, using paired sample t-test.

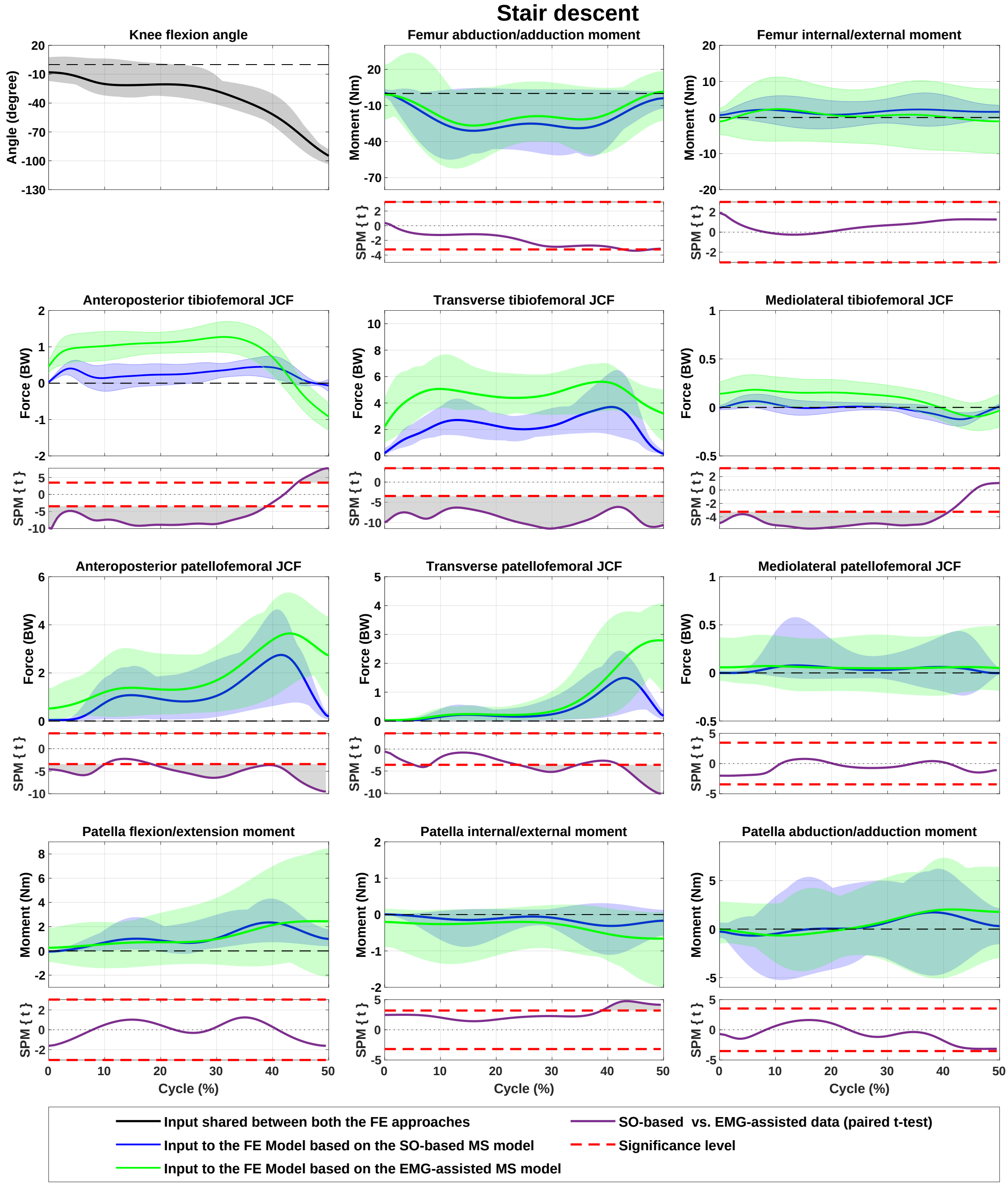


**Fig. S14:** Inputs to the FE models of the study during stair descent, consisted of knee flexion angle, tibiofemoral joint moments (those calculated from inverse dynamics plus the moments generated by the muscles), tibiofemoral joint contact forces, patellofemoral joint contact forces, and patellofemoral moment generated by quadriceps muscles. Inputs estimated by the SO-based and EMG-assisted MS models are illustrated in blue and green, respectively. The SPM plots (in magenta) show comparisons between the results from the EMG-assisted and SO-based MS models, using paired sample t-test.

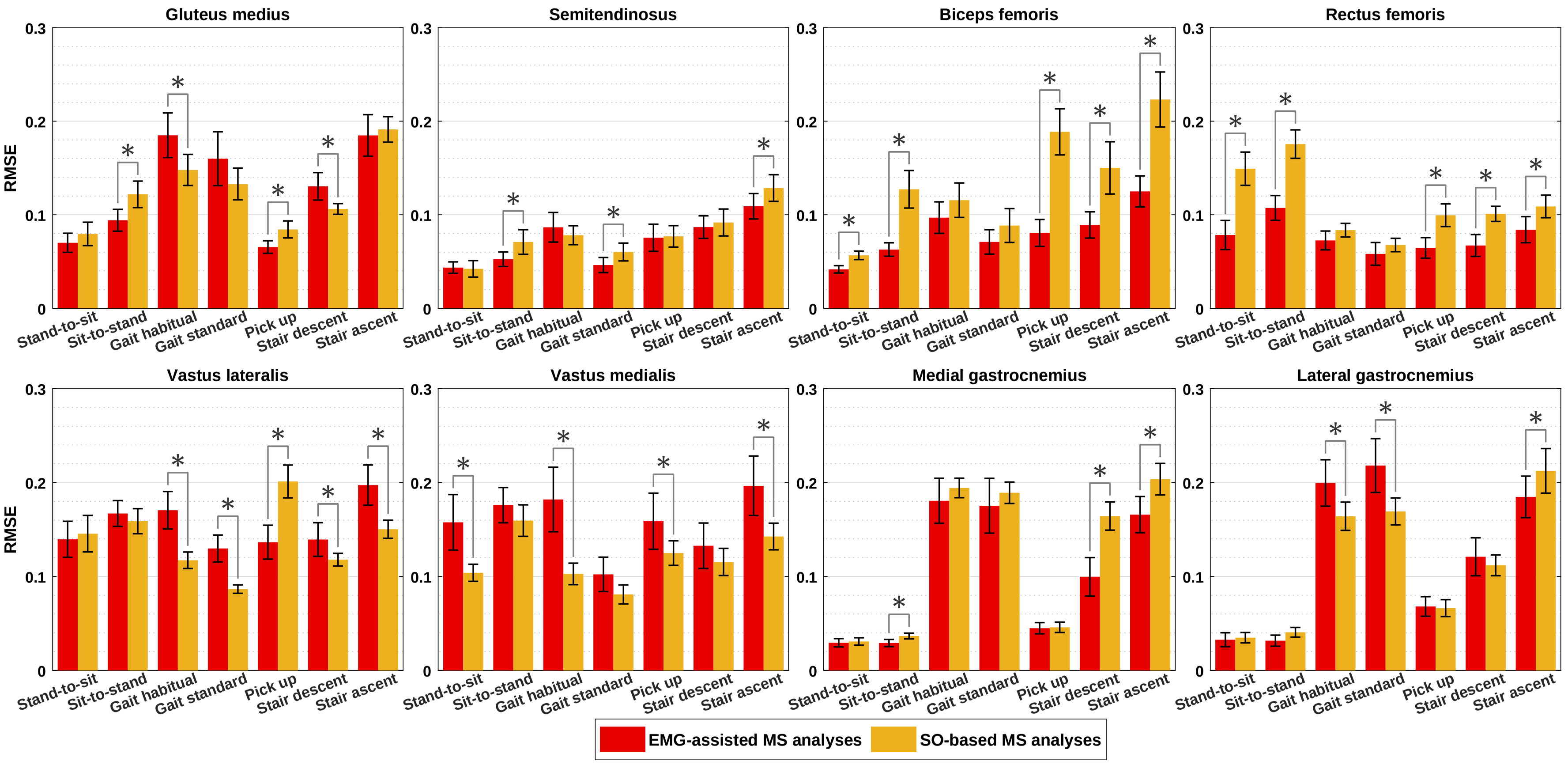


**Fig. S15:** The root mean square error (RMSE) between the enveloped EMGs and estimated muscle activations by the EMG-assisted MS model (in red) and SO-based MS model (in yellow). Stars indicated significant differences (p<0.05) using paired sample t-test, and error bars show 95% confidence intervals.

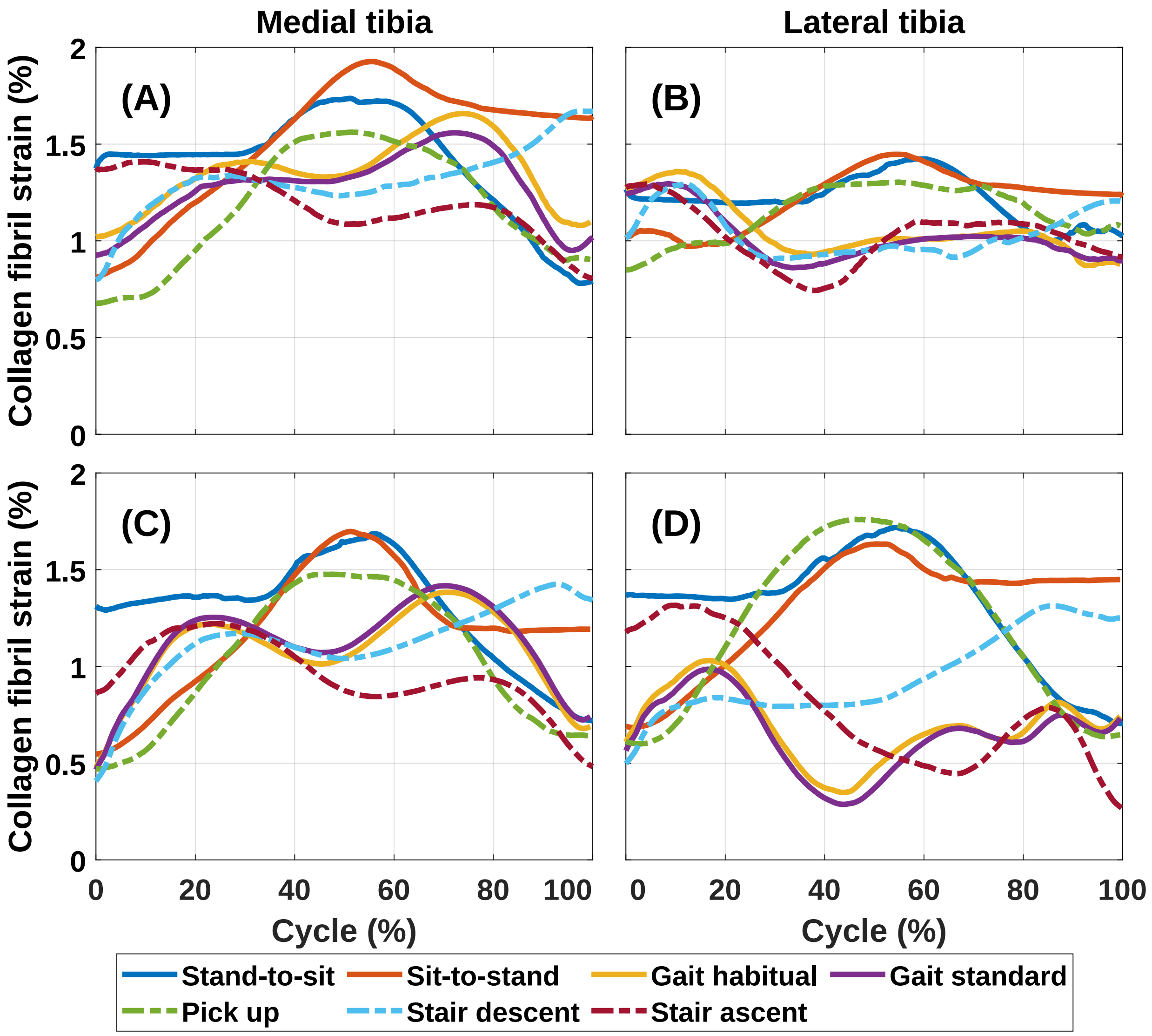


**Fig. S16:** Average of the collagen fibril strain estimated by the EMG-assisted MS-FE models on the medial tibia (A) and lateral tibia (B), and the SO-based MS-FE model on the medial tibia (C) and lateral tibia (D), reporting the 15 subject average profile for each activity. Deviations from the average (e.g., standard deviations) are not shown to improve the readability.

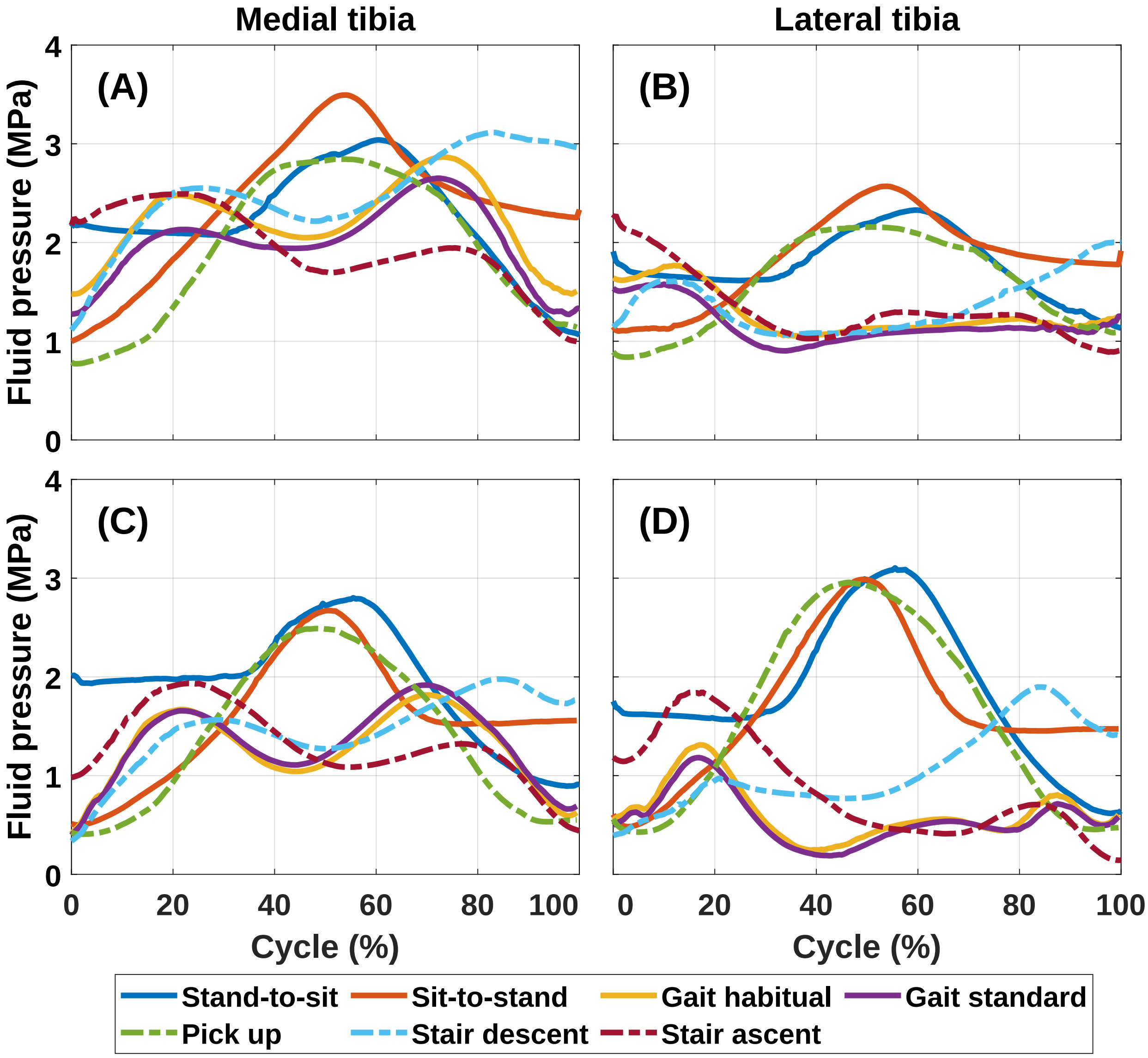


**Fig. S17:** Average of the fluid pressure estimated by the EMG-assisted MS-FE models on the medial tibia (A) and lateral tibia (B), and the SO-based MS-FE model on the medial tibia (C) and lateral tibia (D), reporting the 15 subject average profile for each activity. Deviations from the average (e.g., standard deviations) are not shown to improve the readability.

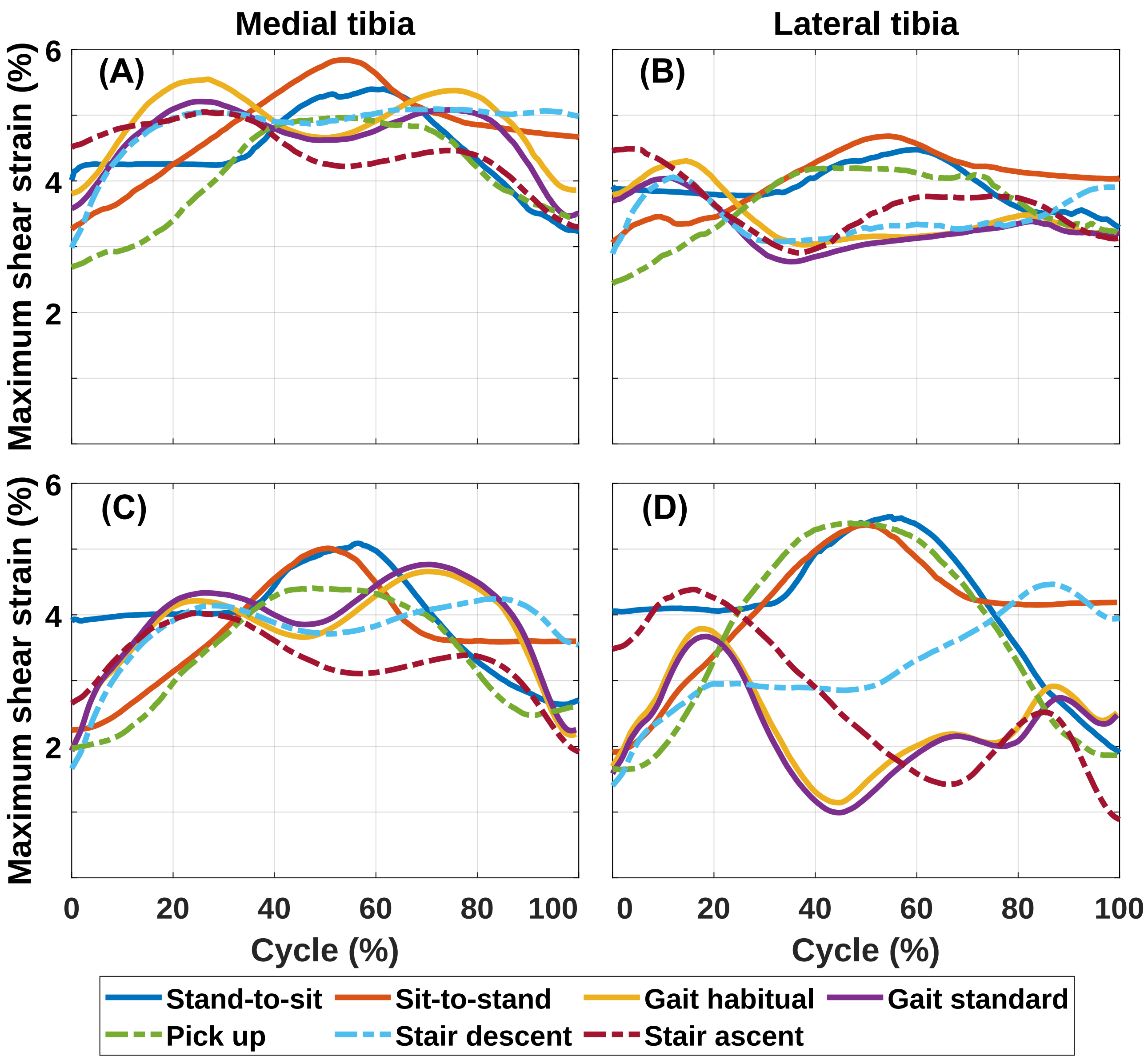


**Fig. S18:** Average of the maximum shear strain estimated by the EMG-assisted MS-FE models on the medial tibia (A) and lateral tibia (B), and the SO-based MS-FE model on the medial tibia (C) and lateral tibia (D), reporting the 15 subject average profile for each activity. Deviations from the average (e.g., standard deviations) are not shown to improve the readability.

### References


[1] R. Altman, E. Asch, D. Bloch, G. Bole, D. Borenstein, K. Brandt, W. Christy, T. Cooke, R. Greenwald, M. Hochberg *et al.*, "Development of criteria for the classification and reporting of osteoarthritis: classification of osteoarthritis of the knee," *Arthritis & Rheumatism: Official Journal of the American College of Rheumatology*, vol. 29, no. 8, pp. 1039–1049, 1986.

[2] H. J. Hermens, B. Freriks, R. Merletti, D. Stegeman, J. Blok, G. Rau, C. Disselhorst-Klug, and G. Hägg, "European recommendations for surface electromyography," *Roessingh research and development*, vol. 8, no. 2, pp. 13–54, 1999.

[3] A. Mantoan, C. Pizzolato, M. Sartori, Z. Sawacha, C. Cobelli, and M. Reggiani, "Motonms: A matlab toolbox to process motion data for neuromusculoskeletal modeling and simulation," *Source code for biology and medicine*, vol. 10, no. 1, pp. 1–14, 2015.

[4] D. G. Lloyd and T. F. Besier, "An emg-driven musculoskeletal model to estimate muscle forces and knee joint moments in vivo," *Journal of biomechanics*, vol. 36, no. 6, pp. 765–776, 2003.

[5] P. Julkunen, P. Kiviranta, W. Wilson, J. S. Jurvelin, and R. K. Korhonen, "Characterization of articular cartilage by combining microscopic analysis with a fibril-reinforced finite-element model," *Journal of biomechanics*, vol. 40, no. 8, pp. 1862–1870, 2007.

[6] W. Wilson, C. Van Donkelaar, B. Van Rietbergen, K. Ito, and R. Huiskes, "Stresses in the local collagen network of articular cartilage: a poroviscoelastic fibril-reinforced finite element study," *Journal of biomechanics*, vol. 37, no. 3, pp. 357–366, 2004.

[7] S. Below, S. P. Arnoczky, J. Dodds, C. Kooima, and N. Walter, "The split-line pattern of the distal femur: A consideration in the orientation of autologous cartilage grafts," *Arthroscopy: The Journal of Arthroscopic & Related Surgery*, vol. 18, no. 6, pp. 613–617, 2002.

[8] P. Boettcher, M. Zeissler, J. Maierl, V. Grevel, and G. Oechtering, "Mapping of split-line pattern and cartilage thickness of selected donor and recipient sites for autologous osteochondral transplantation in the canine stifle joint," *Veterinary surgery*, vol. 38, no. 6, pp. 696–704, 2009.

[9] D. W. Goodwin, Y. Z. Wadghiri, H. Zhu, C. J. Vinton, E. D. Smith, and J. F. Dunn, "Macroscopic structure of articular cartilage of the tibial plateau: influence of a characteristic matrix architecture on mri appearance," *American Journal of Roentgenology*, vol. 182, no. 2, pp. 311–318, 2004.

[10] B. M. Leo, M. A. Turner, and D. R. Diduch, "Split-line pattern and histologic analysis of a human osteochondral plug graft," *Arthroscopy: The Journal of Arthroscopic & Related Surgery*, vol. 20, pp. 39–45, 2004.

[11] Y. Dabiri and L. Li, "Influences of the depth-dependent material inhomogeneity of articular cartilage on the fluid pressurization in the human knee," *Medical engineering & physics*, vol. 35, no. 11, pp. 1591–1598, 2013.

[12] K. S. Halonen, M. Mononen, J. Jurvelin, J. Töyräs, A. Kłodowski, J.-P. Kulmala, and R. Korhonen, "Importance of patella, quadriceps forces, and depthwise cartilage structure on knee joint motion and cartilage response during gait," *Journal of biomechanical engineering*, vol. 138, no. 7, 2016.

[13] W. Lai, V. C. Mow, and V. Roth, "Effects of nonlinear strain-dependent permeability and rate of compression on the stress behavior of articular cartilage," *Journal of Biomechanical Engineering*, vol. 103, no. 2, pp. 61–66, 1981.

[14] H. Lipshitz, R. Etheredge 3rd, and M. J. Glimcher, "In vitro wear of articular cartilage." *The Journal of bone and joint surgery. American volume*, vol. 57, no. 4, pp. 527–534, 1975.

[15] W. Wilson, C. Van Donkelaar, B. Van Rietbergen, K. Ito, and R. Huiskes, "Erratum to "stresses in the local collagen network of articular cartilage: a poroviscoelastic fibril-reinforced finite element study"[journal of biomechanics 37 (2004) 357-366] and" a fibril-reinforced poroviscoelastic swelling model for articular cartilage"[journal of biomechanics 38 (2005) 1195–1204]," *Journal of Biomechanics*, vol. 38, no. 10, pp. 2138–2140, 2005.

[16] E. Danso, J. Mäkelä, P. Tanska, M. Mononen, J. Honkanen, J. Jurvelin, J. Töyräs, P. Julkunen, and R. Korhonen, "Characterization of site-specific biomechanical properties of human meniscus—importance of collagen and fluid on mechanical nonlinearities," *Journal of biomechanics*, vol. 48, no. 8, pp. 1499–1507, 2015.

[17] A. Benninghoff, "Form und bau der gelenkknorpel in ihren beziehungen zur funktion," *Zeitschrift für Zellforschung und mikroskopische Anatomie*, vol. 2, no. 5, pp. 783–862, 1925.

[18] A. Esrafilian, L. Stenroth, M. E. Mononen, P. Tanska, S. Van Rossom, D. G. Lloyd, I. Jonkers, and R. K. Korhonen, "12 degrees of freedom muscle force driven fibril-reinforced poroviscoelastic finite element model of the knee joint," *IEEE Transactions on Neural Systems and Rehabilitation Engineering*, vol. 29, pp. 123–133, 2021.

[19] L. Blankevoort and R. Huiskes, "Ligament-bone interaction in a three-dimensional model of the knee," *Journal of biomechanical engineering*, vol. 113, no. 3, pp. 263–269, 1991.

[20] D. L. Butler, M. D. Kay, and D. C. Stouffer, "Comparison of material properties in fascicle-bone units from human patellar tendon and knee ligaments," *Journal of biomechanics*, vol. 19, no. 6, pp. 425–432, 1986.

[21] P. Atkinson, T. Atkinson, C. Huang, and R. Doane, "A comparison of the mechanical and dimensional properties of the human medial and lateral patellofemoral ligaments," in *Proceedings of the 46th Annual Meeting of the Orthopaedic Research Society, Orlando, FL*, 2000.

[22] D. F. Villegas, J. A. Maes, S. D. Magee, and T. L. H. Donahue, "Failure properties and strain distribution analysis of meniscal attachments," *Journal of biomechanics*, vol. 40, no. 12, pp. 2655–2662, 2007.

[23] L. Schatzmann, P. Brunner, and H. Stäubli, "Effect of cyclic preconditioning on the tensile properties of human quadriceps tendons and patellar ligaments," *Knee Surgery, Sports Traumatology, Arthroscopy*, vol. 6, no. 1, pp. S56–S61, 1998.

[24] M. E. Mononen, M. K. Liukkonen, and R. K. Korhonen, "Utilizing atlas-based modeling to predict knee joint cartilage degeneration: data from the osteoarthritis initiative," *Annals of biomedical engineering*, vol. 47, no. 3, pp. 813–825, 2019.

[25] A. Esrafilian, L. Stenroth, M. Mononen, P. Tanska, J. Avela, and R. Korhonen, "emg-assisted muscle force driven finite element model of the knee joint with fibril-reinforced poroelastic cartilages and menisci," *Scientific reports*, vol. 10, no. 1, pp. 1–16, 2020.

[26] R. J. van Arkel, L. Modenese, A. T. Phillips, and J. R. Jeffers, "Hip abduction can prevent posterior edge loading of hip replacements," *Journal of Orthopaedic Research*, vol. 31, no. 8, pp. 1172–1179, 2013.
